# White Adipose Tissue Compositional and Transcriptional Patterns with Progressive Glucose Intolerance in Persons with HIV

**DOI:** 10.1101/2022.03.21.484794

**Authors:** Samuel S. Bailin, Jonathan A. Kropski, Rama D. Gangula, LaToya Hannah, Joshua D. Simmons, Mona Mashayekhi, Fei Ye, Run Fan, Simon Mallal, Christian M. Warren, Spyros A. Kalams, Curtis L. Gabriel, Celestine N. Wanjalla, John R. Koethe

## Abstract

Subcutaneous adipose tissue (SAT) is a critical regulator of systemic metabolic homeostasis. Persons with HIV (PWH) have an increased risk of metabolic diseases and significant alterations in the SAT immune environment compared with the general population. We generated a comprehensive SAT atlas to characterize cellular compositional and transcriptional changes in 59 PWH with a spectrum of metabolic health. Glucose intolerance was associated with increased lipid-associated macrophages and CD4^+^ and CD8^+^ T effector memory cells, and decreased perivascular macrophages. We observed a coordinated intercellular regulatory program which enriched for genes related to inflammation and lipid-processing across multiple cell types as glucose intolerance increased. Increased CD4^+^ effector memory tissue resident cells most strongly associated with altered expression of adipocyte genes critical for lipid metabolism and cellular regulation. Many of these findings were present in a separate group of 32 diabetic HIV-negative persons, suggesting these changes are not specific to HIV.

## Introduction

The rising global rates of obesity and type 2 diabetes mellitus (T2DM) represent a significant public health challenge^1, 2^. Persons with HIV (PWH) suffer disproportionately from cardiovascular disease, chronic kidney disease, and T2DM compared with the general population^3–6^. The higher risk of metabolic disease in PWH compared with the general population is likely multifactorial and stems from increasing age and survival with improved anti-retroviral therapies, a greater prevalence of known risk factors for metabolic disease including elevated body mass index (BMI), and HIV-specific risk factors including persistent inflammation^3, 7–9^. Adipose tissue is an important regulator of glucose and lipid metabolism^10^, and alterations in its cellular composition and function have been implicated in the development of metabolic diseases^11–15^. While alterations in adipose tissue are common in PWH^7^, the underlying changes in subcutaneous adipose tissue (SAT) that contribute to metabolic disease in this population are poorly understood.

Despite the success of modern antiretroviral therapy (ART), virologically suppressed PWH have persistent innate and adaptive immune activation^16^, as well as permanent alterations in the adipose tissue immune cell compartment^17^. In non-obese PWH, CD8^+^ T cells accumulate in SAT in a process strikingly similar to that observed in obesity in the general population^18, 19^. Additionally, SAT CD4^+^ T cells in PWH shift towards an increased proportion of inflammatory and cytotoxic cells compared to HIV-negative individuals^20^. Many immune cell populations implicated in the perpetuation of chronic SAT inflammation in studies of HIV-negative individuals or animals are also altered in circulating immune cells in PWH^21, 22^. Therefore, HIV may serve as a natural experimental model, to study exaggerated immune activation and cellular changes, which can shed light on SAT pathophysiology shared with the general population. Recent studies in lean and obese persons in the general population have begun to unravel the complex cellular composition of adipose tissue^14, 15, 23, 24^. However, a comprehensive assessment of the identity, polarization, and molecular programs of adipose tissue immune cells in PWH, a group at particularly high risk of metabolic disease, has not been reported.

We recruited a large cohort of PWH with a spectrum of metabolic health to elucidate the SAT cellular compositional and transcriptional regulatory framework that define metabolic disease. We used single-cell proteogenomics and generated a detailed molecular atlas of SAT from 59 PWH to characterize disease-specific changes in cellular populations and expression programs. In PWH with glucose intolerance, we found higher proportion of lipid-associated macrophage (LAM) and LAM-like macrophage populations, reduction of perivascular macrophages (PVMs), and an increase in CD4^+^ and CD8^+^ T effector memory (T_EM_) populations that was independent of obesity, age, and sex. We further uncovered a multicellular transcriptional regulatory program with glucose intolerance characterized by a shift towards macrophage lipid processing phenotype, increasing cytotoxicity and IFN-γ phenotype in T cells, and genes associated with fibrosis. Finally, we showed that this regulatory pattern is conserved in diabetic HIV-negative individuals, suggesting that polarization of immune cells regulating adipose tissue inflammation is a shared feature of metabolic disease and not specific to PWH. These transcriptomic and other data are posted to a user-friendly interactive website (http://vimrg.app.vumc.org/).

## Results

### Cellular Composition Reflects the Complex Function of Subcutaneous Adipose Tissue

We enrolled individuals with a range of glucose intolerance to investigate the role of adipose tissue immune cells in the development of metabolic disease among PWH registered at ClinicalTrials.gov (NCT04451980) (Table 1). To determine the cell populations that characterize SAT in the context of glucose intolerance and HIV infection, we collected abdominal SAT from non-diabetic (n = 20), pre-diabetic (n = 19), and diabetic PWH (n = 20). We used the droplet-based 10X Genomics platform with cellular indexing of transcriptomes and epitopes by sequencing (CITE-seq)^25^, to analyze surface phenotypes and transcriptomes (Fig.1a, Supplementary Fig. 1a, methods). After quality control filtering and harmony batch correction^26^, (Supplementary Fig. 1b-1e, Supplementary Tables 1-3), we obtained a final dataset with 177,942 cells from 59 participants (Fig. 1b, 1c). Broadly, cells clustered into fibro-adipogenic, lymphoid, vascular, and myeloid clusters (Fig. 1d). Cells were annotated using genes previously identified in single-cell datasets (Supplementary Fig. 1f, Supplementary Tables 4 and 5)^15, 27–30^. Importantly, we identified several immune cell types that have been associated with diabetes and obesity in prior studies, including LAMs^28, 31^, natural killer (NK) cells^32^, gamma delta (γδ) T cells^33^, and innate lymphoid cells (ILCs)^15, 34^. We also identified three clusters (arterial EndoMT-like, venous EndoMT-like, and capillary EndoMT-like) expressing markers of both pericytes and endothelial cells^35^. Prediabetic participants had a greater proportion of vascular cells (p_adj_ = 0.03) and reduced proportion of myeloid and lymphoid cells (p_adj_ = 0.04, p_adj_ = 0.02, respectively) compared with non-diabetic participants (Fig. 1e).

**Figure 1.**
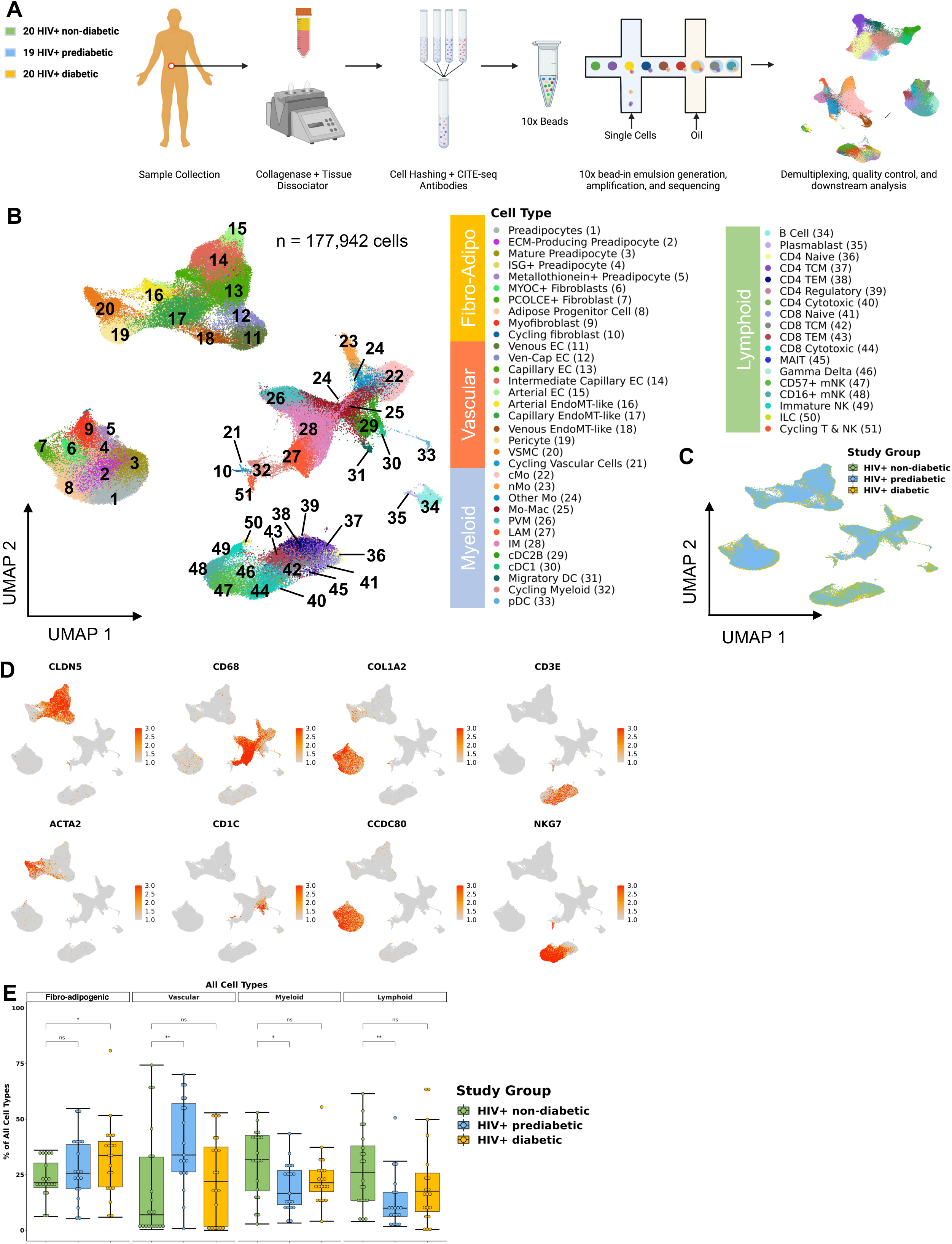
Single-cell RNA sequencing reveals complex cellular structure of subcutaneous white adipose tissue. **a**, Schematic overview of study design. Twenty HIV+ non-diabetic, 19 HIV+ prediabetic, and 20 HIV+ diabetic participants underwent abdominal subcutaneous adipose tissue harvesting with liposuction. Tissue was processed with collagenase and dissociated. Single-cell suspensions from each participant were hashed and labeled with CITE-seq antibodies before multiplexing in groups of four. 10x libraries were generated using the Chromium platform and sequenced on Illumina NovaSeq 6000. The bioinformatic pipeline included demultiplexing, quality control, dimensional reduction and clustering, and transcriptional analysis. **b**, Uniform Manifold Approximation and Projection (UMAP) of 177,942 cells from 59 individuals after removal of doublets and quality control, with manual annotation of cell clusters based on canonical gene markers. **c**, UMAP after harmony integration grouped by disease status showing successful integration; HIV+ non-diabetic (green), HIV+ prediabetic (blue), and HIV+ diabetic (yellow). **d**, Gene expression projected onto the UMAP identifying major cell types including fibro-adipogenic (*COL1A2, CCDC80*), vascular (*CLDN5, ACTA2*), myeloid (CD68, CD1C), and T cell and natural killer cells (*CD3E, NKG7*). **e**, Boxplot showing the proportion of major cell categories (fibro-adipogenic, vascular, lymphoid, and myeloid) as a percentage of total cells split by disease status (n = 59) (HIV+ non-diabetic, green; HIV+ prediabetic, blue; HIV+ diabetic, yellow). The horizontal black line represents the median, the box shows the lower and upper quartile limits and the whiskers are 1.5x the interquartile range. Abbreviations: cMo, classical monocyte; cDC1, conventional dendritic cell type 1; cDC2B, conventional dendritic cell type 2B; DC, dendritic cell; EC, endothelial cell; ECM, extracellular matrix; EndoMT, endothelial-mesenchymal transitional; ILC, innate lymphoid cell; IM, intermediate macrophage; ISG+, interferon; LAM, lipid-associated macrophage; Mac, macrophage; mNK, mature natural killer; Mo, monocyte; NK, natural killer; nMo, non-classical monocyte; pDC, plasmacytoid dendritic cell; PVM, perivascular macrophage; VSMC, vascular smooth muscle cells

**Table 1.** Cohort Characteristics. Continuous variables are shown as median value with interquartile range. Categorical variables are shown as number and percent.

|  | **HIV+ non-diabetic**<br>**(n = 20)** | **HIV+ prediabetic**<br>**(n = 19)** | **HIV+ diabetic**<br>**(n = 20)** | **P value** |
| --- | --- | --- | --- | --- |
| Age, years | 46 (40, 52) | 42 (38, 54) | 55 (48, 62) | 0.053 |
| Race, black (%) | 7 (35) | 8 (42) | 23 (39) | 0.99 |
| Sex, female | 7 (35) | 4 (21) | 5 (25) | 0.60 |
| Body mass index (kg/m<sup>2</sup>) | 31.0 (28.6, 34.7) | 29.4 (28.4, 34.2) | 33.2 (30.2, 37.8) | 0.079 |
| Fasting blood glucose (mg/dl) | 88 (82, 92) | 111 (102, 122) | 184 (120, 242) | < 0.001 |
| Hemoglobin A1c (%) | 5.3 (5.1, 5.4) | 5.5 (5.1, 5.9) | 7.8 (6.4, 9.5) | < 0.001 |
| Metformin use (%) | 0 (0) | 0 (0) | 12 (60) | < 0.001 |
| CD4 count at enrollment | 784 (660, 941) | 684 (544, 1000) | 893 (724, 1102) | 0.53 |
| Thymidine analogue exposure (%) | 5 (26) | 2 (11) | 4 (22) | 0.45 |
| Integrase-based regimen (%) | 13 (65) | 11 (58) | 11 (55) | 0.80 |

We next subset on major cell populations for finer cell annotations. Adipose tissue macrophages and conventional dendritic cells (cDC) interact with adipocytes and can modulate adipogenesis, insulin sensitivity, and tissue remodeling in the setting of obesity^28, 36–39^, though there are few similar data in PWH^40, 41^. We identified 14 distinct cell populations from 40,585 cells in the myeloid compartment (Fig. 2a) using canonical gene markers (Fig. 2b, Supplementary Table 6). Surface-marker phenotyping by CITE-seq supported our classification of LAMs (CD11C^+^), perivascular macrophages (PVMs) (CD11C^-^), non-classical monocytes (nMo) (CD16^+^), classical monocytes (cMo) (CD14^+^), and DCs (CD1C^+^) (Supplementary Fig. 2a). A subset of macrophages was labeled as intermediate macrophage (IM) and expressed markers of LAMs and PVMs (Supplementary Fig. 2b, Supplementary Table 7). Over-representation analysis demonstrates distinct pathways highlighting functional differences between macrophage subsets (Supplementary Fig. 2c).

**Figure 2.**
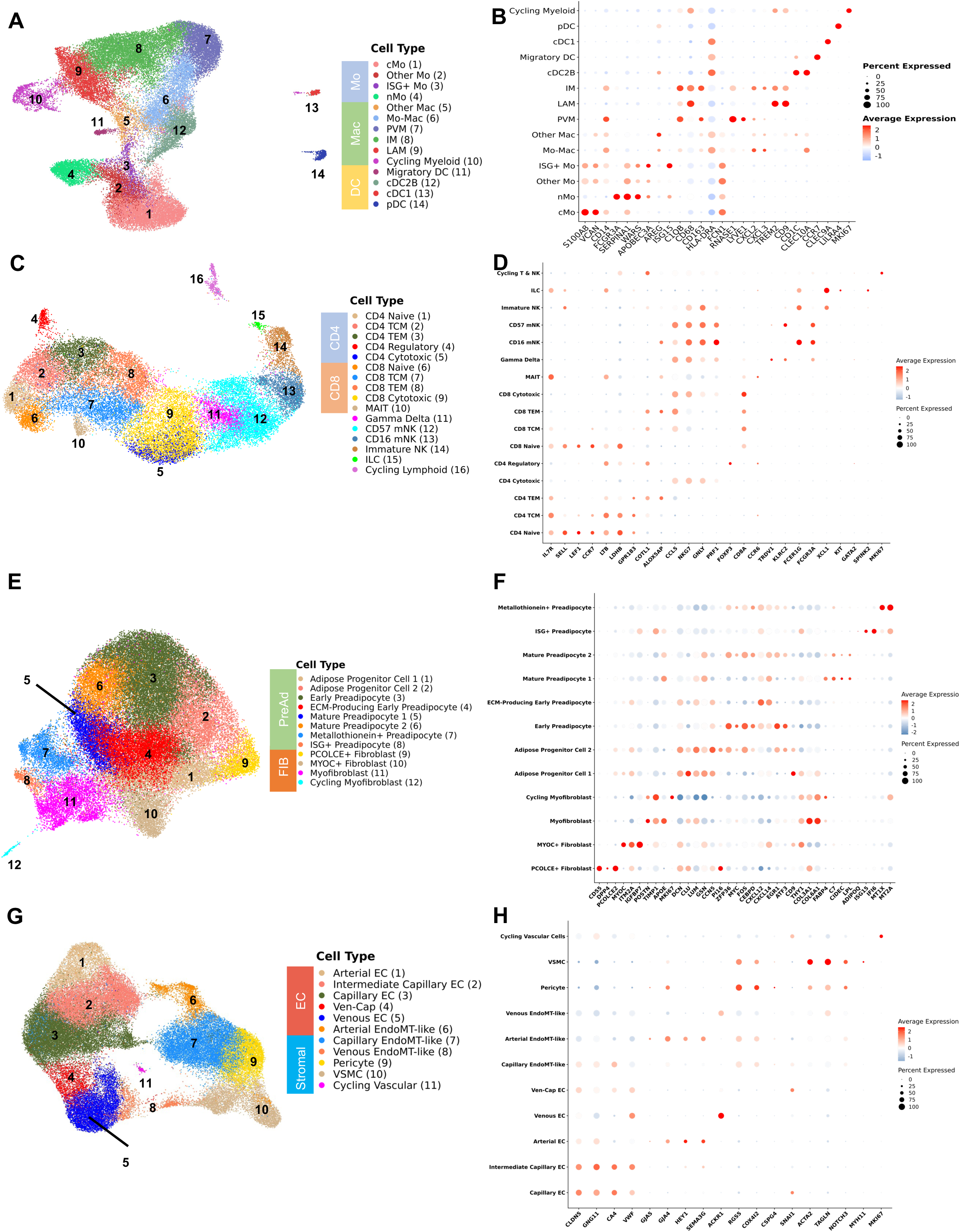
Analysis of Subsets Shows Delineation of Cell Types That are Important for Adipose Tissue Function. **a**, Uniform Manifold Approximation and Projection (UMAP) of myeloid cells (n = 40,585 cells) from 59 individuals after subsetting and reclustering showing 14 distinct cell types/states. **b**, Dot plot showing selected myeloid gene markers on the x-axis and cell type on the y-axis. The dot size reflects the percentage of cells expressing the gene while the color reflects the scaled average expression. **c,** UMAP of T cells, natural killer cells, and innate lymphoid cells (n = 28,061 cells) from 59 individuals after subsetting and reclustering showing 16 distinct cell types/states. **d**, Dot plot showing selected Lymphoid gene markers on the x-axis and cell type on the y-axis. The dot size reflects the percentage of cells expressing the gene while the color reflects the scaled average expression. **e**, UMAP of fibro-adipogenic cells (n = 51,029 cells) from 59 individuals after subsetting and reclustering. **f**, Dot plot showing selected fibro-adipogenic gene markers on the x-axis and cell type on the y-axis. The dot size reflects the percentage of cells expressing the gene while the color reflects the scaled average expression. **g**, UMAP of vascular cells (n = 55,534 cells) from 59 individuals after subsetting and reclustering. **h**, Dot plot showing selected vascular gene markers on the x-axis and cell type on the y-axis. The dot size reflects the percentage of cells expressing the gene while the color reflects the scaled average expression. Abbreviations: cMo, classical monocyte; cDC1, conventional dendritic cell type 1; cDC2B, conventional dendritic cell type 2B; DC, dendritic cell; EC, endothelial cell; ECM, extracellular matrix; EndoMT, endothelial-mesenchymal transitional; ILC, innate lymphoid cell; IM, intermediate macrophage; IM, intermediate macrophage; ISG+, interferon; LAM, lipid-associated macrophage; Mac, macrophage; mNK, mature natural killer; Mo, monocyte; NK, natural killer; nMo, non-classical monocyte; pDC, plasmacytoid dendritic cell; PVM, perivascular macrophage; VSMC, vascular smooth muscle

Lymphoid cells also have a prominent role in shaping the immune environment of SAT and modulating local inflammation and insulin resistance^13, 42–45^. HIV infection induces broad changes to circulating lymphoid cells associated with the development of insulin resistance including memory, senescent, and exhausted phenotypes^19, 21, 22, 46^. From 28,061 lymphoid cells, we identified 16 distinct cell states in adipose tissue including CD4^+^ & CD8^+^ naïve (*SELL, LEF1, CCR7*), central memory (T_CM_) (*LTB, LDHB, GPR183*), effector memory (T_EM_) (*ALOX5AP, COTL1, CCL5*), cytotoxic (*NKG7, GNLY, PRF1*), and CD4^+^ T_reg_ (*FOXP3, CTLA4*) cells. Additionally, we identified γδ T cells (*TRDV1, KLRC2*), MAIT cells (*CCR6, IL7R, KLRG1*), and ILCs *(XCL1, KIT, IL7R*). Several NK cell subsets were identifiable by cell-surface expression of CD16 and CD56 and included mature CD16^+^ (*FCER1G, NKG7)*, terminally differentiated mature CD16^+^CD57^+^ (*FCGR3A, KLRD1*), and CD56^+^CD16^-^ (*XCL1, XCL2, SELL*) (Fig. 2c, 2d; Supplementary Table 6). The RNA transcriptome profiles were analyzed in parallel with CITE-seq surface marker expression of CD4, CD8, CD45RA, CD27, CD57, and CD16 (Supplementary Fig. 2d). We also separately assessed only CD4^+^ and CD8^+^ T cells to differentiate more finely between naïve, central memory, effector memory, and cytotoxic phenotypes (Supplementary Fig. 2E-2H, Supplementary Table 7). Cells classified as CD4^+^ T_EM_ expressed higher levels of CD69, a marker of tissue residency (Supplementary Fig. 2f)^47^. We validated this by gating on CD4^+^ CD69^+^ T cells for 21 individuals and found high correlation with CD4^+^ T_EM_ defined by scRNA-seq (ρ = 0.71, p < 0.001). These data suggest that expression of CD69 is a major driver of gene expression pattern.

We next evaluated fibro-adipogenic cell populations, as recent studies have revealed considerable heterogeneity^14, 23, 24, 48–51^ and interactions between fibro-adipogenic and immune cells can modulate adipocyte function and have an important role in the development of metabolic disease^39, 52^. We identified 11 distinct cells states from 51,029 fibro-adipogenic cells (Fig. 2e, 2f; Supplementary Table 6). An interstitial fibroblast population (PCOLCE^+^ fibroblast) that has been shown to give rise to preadipocytes expressing *SFRP2, DPP4,* and *PCOLCE2* is analogous to PCOLCE^+^ fibroblasts in the dermis^53^, and homologous to a population in mice^30^. A second fibroblast population has high expression of *MYOC* and *ITM2A*, which is consistent with anti-adipogenic CD142 cells that have been previously described^48^. A separate population of cells expressing *TIMP1* and *POSTN* is transcriptionally consistent with myofibroblasts. Adipose progenitor cells (PCs) were characterized by expression of *DCN*, *CLU, LUM, GSN,* and *PI16*. The preadipocyte compartment was largely differentiated by expression immediate early genes (*MYC, FOS, JUN*) and markers associated with adipogenesis and regulation of inflammation (*ZFP36, EGR1, KLF4, CEBPD*) (Early preadipocyte), and markers associated with extracellular matrix (ECM) (ECM-Producing preadipocytes). Preadipocytes progressively acquired higher expression of *FABP4, LPL,* and *CIDEC*, which are markers of lipid acquisition. Finally, we found different vascular stromal and endothelial cells present in adipose tissue (Fig. 2g, 2h). In summary, we show a diversity of cell types present in SAT in persons with HIV that reflect the complex physiologic functions of adipose tissue.

### Categorical and Continuous Measures of Glucose Intolerance are Associated with Macrophage and T Cell Polarization

Cellular compositional changes of SAT with progressive glucose intolerance likely reflect functional changes that either promote or are derived from the disease process. Several studies evaluating obesity have shown significant changes to the myeloid compartment with increasing proportion of LAMs and changes to T cell polarization. Less is known about compositional changes to the fibro-adipogenic compartment, though it is known that adipose tissue fibrosis occurs in the setting of obesity^54^.

Within the macrophage compartment, IM and LAM proportions were higher in prediabetic (p_adj_ = 0.02 and p_adj_ = 0.03) and diabetic (p_adj_ = 0.02 and p_adj_ = 0.04) PWH compared with non-diabetic PWH. In contrast, PVM proportion was lower in prediabetic (p_adj_ = 0.02) and diabetic PWH (p_adj_ = 0.02) compared with non-diabetic PWH (Fig. 3a). In general, prediabetic and diabetic PWH had a shift away from classical monocyte (cMo) (p_adj_ = 0.02 & p_adj_ = 0.09, respectively) and other Mo (p_adj_ = 0.02 & p_adj_ = 0.02, respectively). Compared with non-diabetic PWH, cDC1 proportion was increased in diabetic PWH (p_adj_ = 0.04). A major strength of this large dataset over previous adipose tissue single cell studies is greater power to assess the independent contributions of important biological factors to cellular composition. We examined whether the cell types associated with glucose intolerance in group comparisons were associated with measurements of glucose intolerance (hemoglobin A1c [hba1c], fasting blood glucose [FBG]) as a continuous measure. We used partial spearman’s correlation adjusted for BMI, sex, and age, and we excluded diabetic PWH from the analysis given the effects of medication treatment on the endpoints. FBG was inversely associated with PVM proportion (ρ = -0.61, p = 0.002) while HbA1c was associated with LAM proportion (ρ = 0.47, p = 0.01) (Fig. 3b). Evaluating all myeloid cells, measures of glucose intolerance were significantly associated with LAM and IM proportions and inversely associated with monocyte cell proportions, particularly with HbA1c (Fig. 3c). Thus, there is an overall shift from primarily monocytes and PVMs towards a LAM-like phenotype with glucose intolerance, similar to findings in HIV-negative persons^15^.

**Figure 3.**
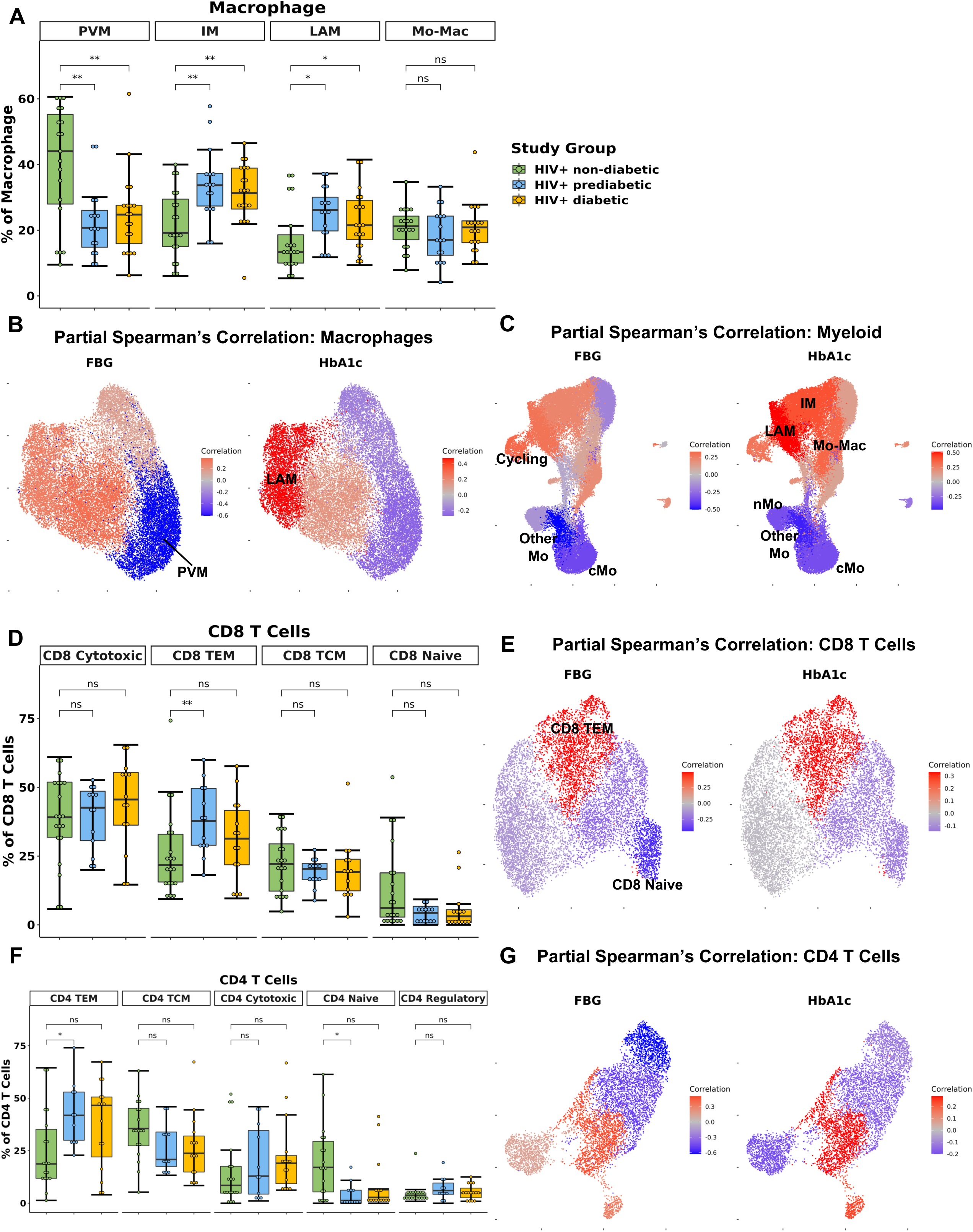
Lipid-associated, intermediate macrophages, and CD4^+^ and CD8^+^ T effector memory proportions are associated with glucose intolerance. **a**, Boxplot showing the proportion of macrophage types split by disease state (HIV+ non-diabetic, green; HIV+ prediabetic, blue; HIV+ diabetic, yellow) (n = 54). The horizontal black line represents the median, the box shows the lower and upper quartile limits and the whiskers are 1.5x the interquartile range. b, Partial spearman’s correlations between fasting blood glucose (FBG) or hemoglobin A1c (HbA1c) and macrophage cell proportions. Spearman’s r for the biological factor (FBG or HbA1c) and each cluster proportion was calculated. The r value (blue, negative; red, positive) for each cluster was plotted onto the macrophage uniform manifold approximation and projection (UMAP). Clusters with significant correlation (p < 0.05) are labeled. c, Partial spearman’s correlation between FBG or HbA1c and cluster proportion plotted onto the myeloid UMAP. d, Boxplot showing the proportion of CD8^+^ T cell subsets as a proportion of total CD8^+^ T cells (n = 47). e, Partial spearman’s correlation between FBG or HbA1c and cluster proportion plotted onto the CD8^+^ T cell UMAP. f, Boxplot showing the proportion of CD4^+^ T cell subsets as a proportion of total CD4^+^ T cells (n = 44). g, Partial spearman’s correlation between FBG or HbA1c and cluster proportion plotted onto the CD4^+^ T cell UMAP. Abbreviations: IM, intermediate macrophage; LAM, lipid-associated macrophage; Mo-Mac, monocyte-macrophage; PVM, perivascular macrophage; TCM, T central memory; TEM, T effector memory

The lymphoid compartment had fewer differences by diabetes status. The proportion of CD8^+^ T_EM_ was lower in non-diabetic PWH compared with prediabetic (p_adj_ = 0.05) but not diabetic PWH (p_adj_ = 0.44) (Fig. 3d). FBG but not HbA1c was significantly associated with CD8^+^ T_EM_ (ρ = 0.49, p < 0.001) and inversely associated with CD8^+^ naïve T cells (ρ = -0.44, p = 0.01) (Fig. 3e). Similarly, the proportion of CD4^+^ T_EM_ was lower in non-diabetic PWH compared with prediabetic (p_adj_ = 0.05) but not diabetic PWH (Fig. 3f). This was mainly due to a significant decrease in CD4^+^ naive T cells (p_adj_ = 0.05). As discussed in the previous section, CD4^+^ T_EM_ cells expressed CD69. We confirmed with flow cytometry that participants with glucose intolerance had higher proportion of CD4^+^ CD69^+^ T cells (median percent: non-diabetic (6.4%), prediabetic (37.5%), diabetic (37.7%); p = 0.01). FBG was significantly associated with CD4^+^ T_EM_ (ρ = 0.51, p < 0.001) and inversely associated with CD4^+^ naïve T cells (ρ = -0.63, p < 0.001) (Fig. 3g). The proportion of γδ T cell was increased in prediabetic (p = 0.04) but not diabetic PWH (p = 0.54), though was not significant after correction for multiple comparisons. FBG was independently associated with γδ T cell (ρ = 0.33, p = 0.04) and CD57^+^ mNK (ρ = 0.35, p = 0.05) proportion, but hbA1c was not (Supplementary Fig. 3a). In summary, prediabetic, but not treated diabetic PWH, have increased T_EM_ cells compared with non-diabetic PWH.

Finally, we hypothesized that the proportion of fibroblast populations would be increased with glucose intolerance. However, there was no significant associations between the proportion of fibro-adipogenic cells and continuous measures of glucose intolerance (Supplementary Fig. 3b) or by diabetes status (Supplementary Fig. 3c). While diabetic PWH had a greater proportion of venous EndoMT-like cells (p = 0.008) and intermediate capillary endothelial cells (p = 0.02), there were no significant differences after correction for multiple comparison and no association with measures of glucose intolerance (Supplementary Fig. 3d). Taken together, we show that in the context of glucose intolerance, PWH have dramatic changes in the myeloid cell compartment with a shift towards a LAM-like phenotype, as well as a shift towards T_EM_ T cells, but no significant difference in the fibro-adipogenic cell composition.

### BMI is Associated with Macrophage Polarization while Sex is Associated with Fibro-Adipogenic Compositional Changes

Demographic variables can influence the adipose tissue environment^10^. Therefore, we next examined the relationship of cell composition with BMI, age, and sex. Evaluating all myeloid cells, BMI was associated with the proportion of IMs (ρ = 0.35, p = 0.01), LAMs (ρ = 0.27, p = 0.05), and Mo-Macs (ρ = 0.31, p = 0.03) (Fig. 4a). Similarly, age was associated with the proportion of LAMs (ρ = 0.31, p = 0.03). Evaluating the CD4^+^ T cell subsets, age was inversely related to naïve (ρ = -0.34, p = 0.04) and regulatory T cells (ρ = -0.41, p = 0.02) (Fig. 4b). BMI was inversely related to CD8^+^ T_CM_ proportion (ρ = -0.38, p = 0.01) (Fig. 4c). Finally, age was inversely associated with the proportion of Progenitor 1 (ρ = -0.31, p = 0.03) (Fig. 4d). Other lymphoid and vascular cell proportions were associated with BMI and age (Supplementary Fig. 4a, 4b). In summary, higher BMI and to a lesser extent older age, was significantly associated with a shift towards LAM and LAM-like macrophages.

**Figure 4.**
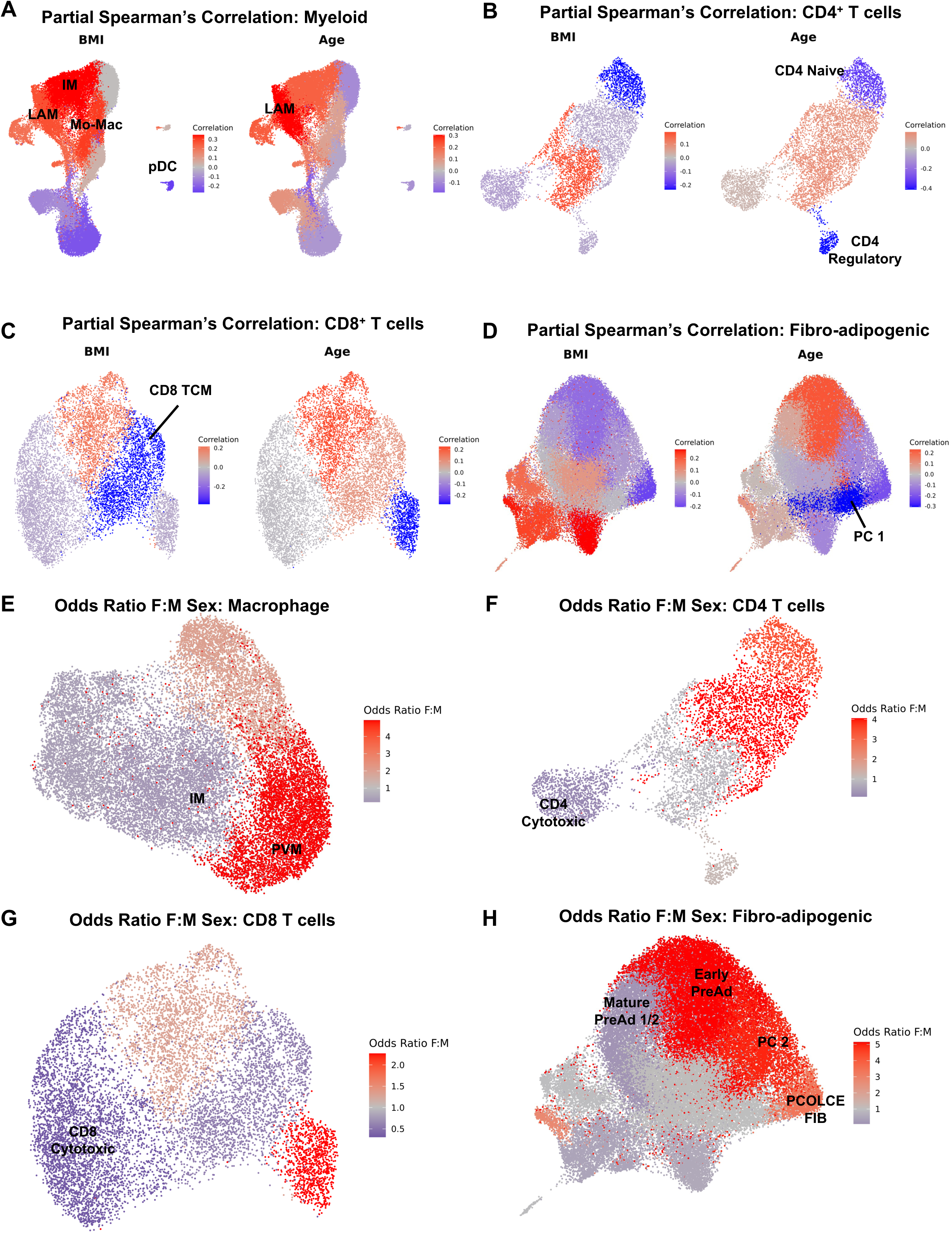
Body Mass Index and Sex Are Associated with Compositional Changes in Immune and Fibro-adipogenic Cells. **a-d**, Partial spearman’s correlations. Spearman’s r for the biological factor (body mass index [BMI] or age) and each cluster proportion was calculated. The r value (blue, negative; red, positive) for each cluster was plotted onto the uniform manifold approximation and projection (UMAP). Clusters with significant correlation (p < 0.05) are labeled. BMI and age plotted onto **a,** Myeloid UMAP, **b,** CD4^+^ T cell UMAP, **c**, CD8^+^ T cell UMAP, and **d,** Fibro-adipogenic UMAP. **e-h,** Ordinal linear regression with cluster proportion as the outcome and sex as the independent variable adjusted for age, BMI, and diabetes status. The regression coefficient for sex was converted into an odds ratio (female:male) and plotted per cluster onto the UMAP. Odds ratio female: male plotted onto **e**, Macrophage UMAP, **f**, CD4^+^ T cell UMAP, **g,** CD8^+^ T cell UMAP, and **h,** fibro-adipogenic UMAP. Abbreviations: PC, progenitor cell; BMI, body mass index; FIB, fibroblast; IM, intermediate macrophage; Mac, macrophage; Mo, monocyte; LAM, lipid-associated macrophage; pDC, plasmacytoid dendritic cell; PreAd, preadipocyte; PVM, perivascular macrophage; TCM, T central memory

To assess the independent contributions of sex to cell proportions, we used an ordinal linear regression model adjusted for age, BMI, and diabetes status. Compared with men, women had higher proportion of PVMs (p = 0.008) and lower proportion of IMs (p = 0.008), but no significant differences in other myeloid cells (Fig. 4e and Supplementary Fig. 4c). Compared with men, women had lower proportion of CD4^+^ cytotoxic (p = 0.002) and CD8^+^ cytotoxic T cells (p = 0.05) (Fig. 4f, 4g). Women also had proportion difference in NK cells (Supplementary Fig. 4d). In contrast to BMI and age, female sex had a significant influence on fibro-adipogenic cell composition. Compared with men, women had higher proportion of PCOLCE+ fibroblasts (p = 0.03), PC 2 (p = 0.004), and early preadipocytes (p = 0.003), and lower proportion of mature preadipocytes 1 (p < 0.001) and 2 (p < 0.001) (Fig. 4h). Vascular cell populations were not associated with female sex (Supplementary Fig. 4e). The individual odds ratios (female: male) are available at (http://vimrg.app.vumc.org/). Taken together, sex appears to be a larger driver of compositional changes in the fibro-adipogenic compartment than either BMI or age. Female sex is associated with higher proportions of cell populations that are abundant in healthy adipose tissue.

### Cell Proportions Associated with Glucose Intolerance are Highly Inter-Related

We next wanted to examine the inter-relatedness of cell populations associated with glucose intolerance using partial spearman’s correlation, adjusted for BMI, age, sex, and diabetes status. We hypothesized that immune cell populations associated with glucose intolerance would be related to each other and may be associated with fibrogenic cells in adipose tissue. The proportion of CD4^+^ T_EM_ was significantly associated with the proportion of several myeloid cells including IMs (ρ = 0.55, p = 0.001), LAMs (ρ = 0.48, p = 0.01), PVMs (ρ = 0.48, p = 0.006), and Mo-Macs (ρ = 0.66, p < 0.001) and inversely associated with cMos (ρ = -0.70 p < 0.001) and other Mos (ρ = -0.70, p < 0.001) (Fig. 5a). The proportion of CD8^+^ T_EM_ was also associated with the proportion of several myeloid cells including IMs (ρ = 0.42, p = 0.01), LAMs (ρ = 0.46, p = 0.006), PVMs (ρ = 0.38, p = 0.02), and cDC1s (ρ = 0.45, p = 0.002), and inversely associated with cMos (ρ = -0.57, p < 0.001) and other monocytes (ρ = -0.72, p < 0.001) (Fig. 5b). We next looked at the fibro-adipogenic cells. The proportion of CD4^+^ T_EM_ was associated with the proportion of ECM-producing early preadipocytes (ρ = 0.46, p = 0.005) and myofibroblasts (ρ = 0.35, p = 0.04), and inversely associated with early preadipocytes (ρ = -0.51, p = 0.006)(Fig. 5c). The proportion of CD8^+^ T_EM_ was similarly associated with the proportion of ECM-producing early preadipocytes (ρ = 0.49, p = 0.004) and inversely associated with early preadipocytes (ρ = -0.49, p = 0.001) (Fig. 5d). The proportion of IM was associated with the proportion of myofibroblasts (ρ = 0.77, p < 0.001) and inversely associated with PCOLCE+ fibroblasts (ρ = -0.74, p < 0.001) as well as several other cell types to a lesser degree (Fig. 5e). The proportion of LAMs was associated with myofibroblasts (ρ = 0.33, p = 0.04) and inversely associated with PCOLCE+ fibroblasts (ρ = -0.30, p = 0.02) (Fig. 5f). Finally, the proportion of PVMs was associated with PCOLCE+ fibroblasts (ρ = 0.50, p < 0.001) and inversely associated with myofibroblasts (ρ = -0.59, p < 0.001) (Fig. 5g). Taken together, we find high correlation between the proportion of T_EM_ cells, macrophages, and fibro-adipogenic cell populations in SAT, including several cells that are associated with glucose intolerance, suggesting a coordinated shift in multiple cell lineages accompanying changes in metabolic heath.

**Figure 5.**
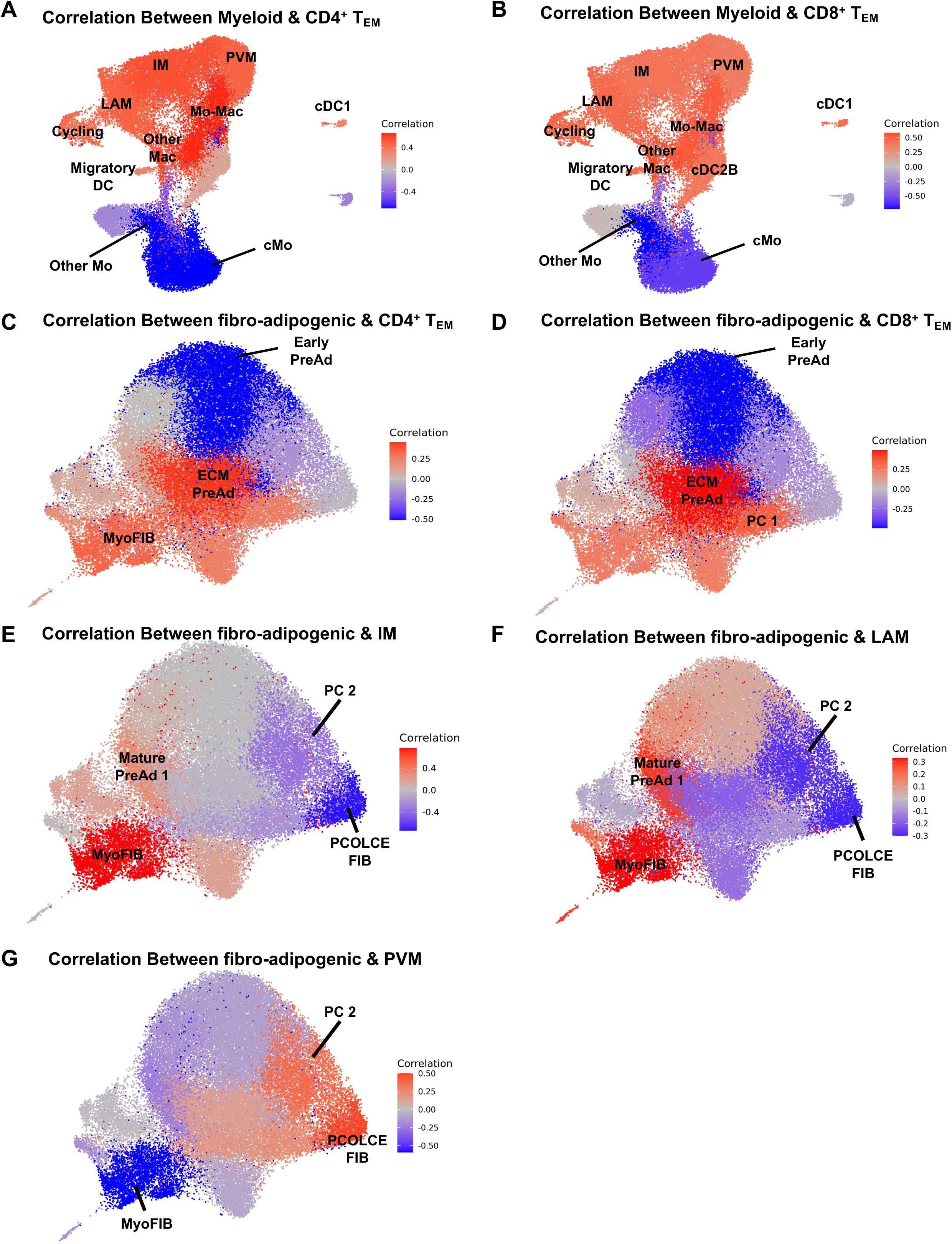
Inter-Cellular Correlation of Proportions with Cells Associated with Glucose Intolerance. **a-g**, Partial spearman’s correlations. Spearman’s r for the immune cell proportion of interest and each cluster proportion was calculated. The r value (blue, negative; red, positive) for each cluster was plotted onto the uniform manifold approximation and projection (UMAP). Clusters with significant correlation (p < 0.05) are labeled. **a,** Spearman’s r for CD4^+^ T effector memory (T_EM_) proportion and myeloid clusters plotted onto the myeloid uniform manifold approximation and projection (UMAP). **b,** Spearman’s r for CD8^+^ T_EM_ cell proportion and myeloid clusters plotted onto the myeloid UMAP. **c**, Spearman’s r for CD4^+^ T_EM_ proportion and fibro-adipogenic clusters plotted onto the fibro-adipogenic UMAP. **d,** Spearman’s r for CD8^+^ T_EM_ proportion and fibro-adipogenic clusters plotted onto the fibro-adipogenic UMAP. **e**, Spearman’s r for intermediate macrophage proportion and fibro-adipogenic clusters plotted onto the fibro-adipogenic UMAP. **f**, Spearman’s r for lipid-associated macrophage proportion and fibro-adipogenic clusters plotted onto the fibro-adipogenic UMAP. **g,** Spearman’s r for perivascular macrophage proportion and fibro-adipogenic clusters plotted onto the fibro-adipogenic UMAP. Abbreviations: BMI, body mass index; cDC1, conventional dendritic cell type 1; cMo, classical monocyte; ECM, extracellular matrix; FIB, fibroblast; IM, intermediate macrophage; Mac, macrophage; Mo, monocyte; MyoFIB, myofibroblast; LAM, lipid-associated macrophage; PC, progenitor cell; pDC, plasmacytoid dendritic cell; PreAd, preadipocyte; PVM, perivascular macrophage.

### Transcriptional Polarization of Macrophages and T Cells towards LAM-like and Effector Memory Phenotype

Having demonstrated that glucose intolerance is independently associated with accumulation of IM, LAM, CD4^+^ T_EM_, and CD8^+^ T_EM_, and inversely associated with PVM, we next examined the transcriptome that defines these cell compartments. We aggregated gene counts for each participant (pseudobulk method) to evaluate the differential gene expression between prediabetic and non-diabetic PWH using a negative binomial generalized linear model implemented in DESeq2^55^, adjusting for age, sex and BMI. 644 genes were differentially expressed (p_adj_ < 0.05) between macrophages from prediabetic and non-diabetic PWH. Macrophages from prediabetics had higher expression of genes related to lipid processing (*FABP5, MGLL, C3, OXSM, SCD, GSTZ1, DBI, ACOT8, DECR1, ETFB*) and oxidative phosphorylation (*NDUFS5, NDUFAB1, ATP5MC3, UQCR10, COX5B*), and lower expression of genes related to chemotaxis (*CCL8, CXCL1, CCL2, CCL4L2, CCL4, CCL3L1, CXCL8, CCL3,*), TNF inflammatory pathway (*TNF, TNFSF18, TNFRSF1A, TNFRSF21*), and M2-macrophage polarization (*TRIB1, EGR1, EGR2, MRC1, LYVE1, KLF4*) (Fig. 6a, Supplementary Table 8). KEGG gene set enrichment analysis (GSEA) confirmed enrichment of metabolic processing pathways and reduction in pathways related to cytokine-cytokine receptor interaction, chemokine signaling, and IL-17 signaling (Fig. 7b). Individual macrophage subsets had fewer transcriptional differences. PVMs from prediabetic PWH had higher expression of *ID3*, and lower expression of *CCL2* compared with non-diabetic PWH. Several genes related to chemokines and M2-phenotypic genes (*TRIB1, HMOX1, CCL2, CCL4*) trended towards lower expression in prediabetic PWH (p_adj_ < 0.1) (Supplementary Table 8). Pseuodotime using Slingshot^56^, demonstrated a single trajectory of monocyte-macrophages transitioning into PVMs (Supplementary Fig. 5a and 5b). Using Tradeseq,^57^ which fits a negative binomial generalized additive model to each gene, expression of transcription factors associated with M2 phenotype increased along the pseudotime trajectory including *ZNF331, NR4A3, KLF4, KLF2, MAF, MAFB, ATF4, EGR2, CEBPD, JUND,* and *SON* (Fig. 6c). Several of these genes had reduced expression in prediabetic compared with non-diabetic PWH with pseudobulk differential gene expression analysis. Despite their large proportion, IM did not demonstrate any differentially expressed genes, suggesting their accumulation, rather than altered transcriptional regulation, is a major driver of phenotypic differences (Supplementary Table 8). Finally, LAMs in prediabetic PWH had higher expression of inflammatory genes (*MIF, HAMP*) and lipid-processing/metabolic genes (*MTLN*, *BSCL2, COX6A1, NDUFB3*), and lower expression of several M2 transcription factors (*TRIB1, EGR1, KLF2*) (Supplementary Fig. 5c, Supplementary Table 8). CD4^+^ T cells in prediabetic PWH had higher expression of markers of activation and effector memory phenotype (*IL32, BATF, CD63, IFITM3, CSTD*) and lower expression of markers related to naïve T cells (*SELL, LEF1, CCR7*) (Fig. 6d). GSEA demonstrated enrichment of pathways related to secretion and cytoskeleton organization (Fig. 6e). CD8^+^ T cells in prediabetic PWH had higher expression of cytotoxic and activation genes (*GZMA, CD63, CLEC2B*) and a trend towards higher expression of *IFNG* (p_adj_ = 0.09) (Fig. 6f).

**Figure 6.**
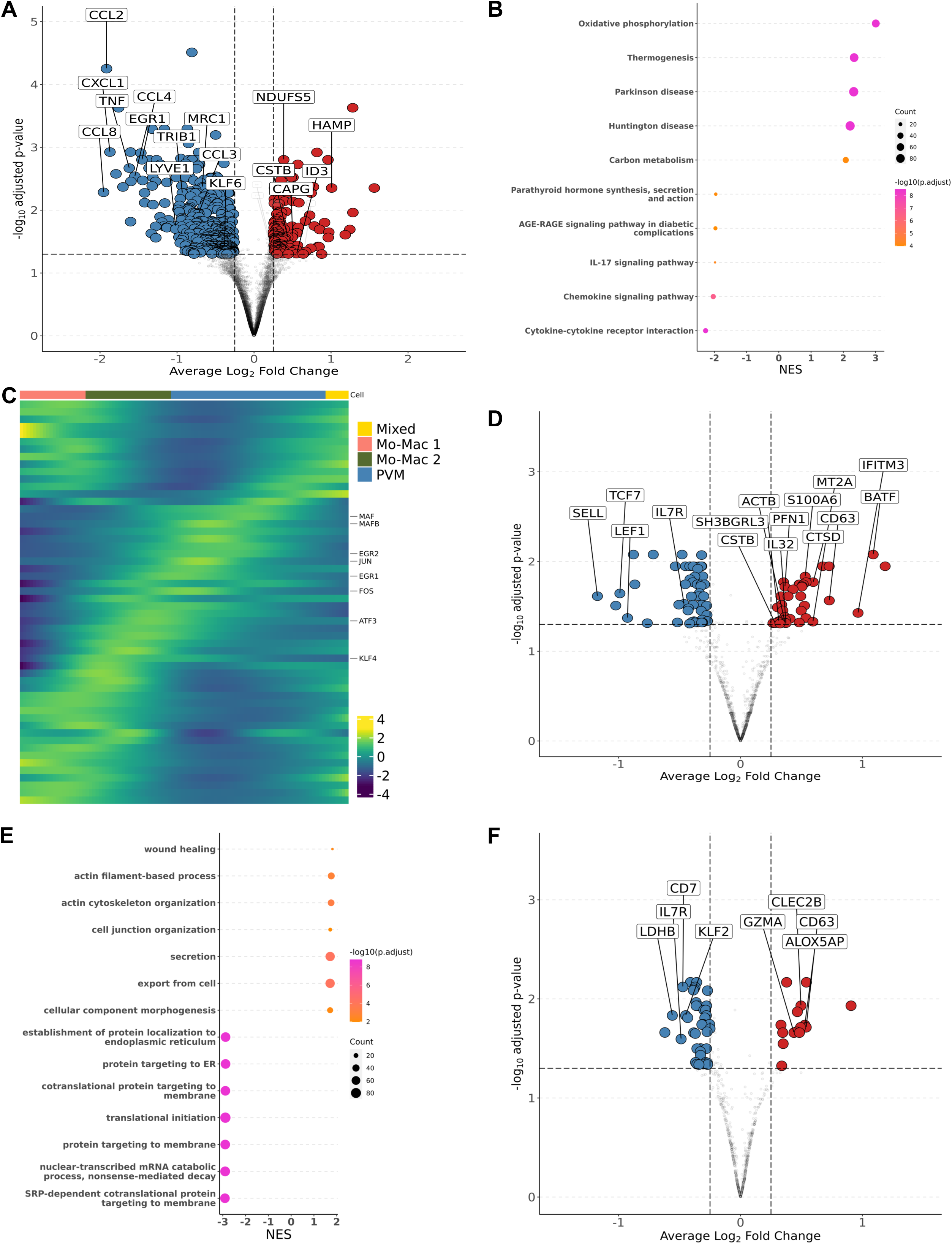
Transcriptional Shift from Immunoregulatory to Metabolic Phenotype in Macrophages and Effector Memory Phenotype in T cells with Glucose Intolerance. **a**, Macrophage volcano plot with average Log_2_ fold change (x-axis) and –log_10_ p-value (y-axis) for prediabetic vs non-diabetic (reference) PWH. Genes that had ≥ 0.25 log_2_ fold change and adjusted p-value < 0.05 were colored red (higher expression) and blue (lower expression). b, Gene set enrichment analysis (GSEA) using the KEGG database. The top 5 and bottom 5 enriched pathways were included with normalized enrichment score (NES) on x-axis and descriptive term on y-axis. Dot size represents the number of gene hits in the pathway and dot color represents the –log_10_ p-value. C, Ordered and smoothed transcription factor gene expression (scaled) along the pseudotime trajectory for monocyte-macrophage to perivascular macrophage. Selected genes were significantly differentially expression along the pseudotime with ≥ log_2_(2) fold change according to TradSeq. d, CD4^+^ T cell volcano plot with average Log_2_ fold change (x-axis) and –log_10_ p-value (y-axis) for prediabetic vs non-diabetic (reference) PWH. Genes that had ≥ 0.25 log_2_ fold change and adjusted p-value < 0.05 were colored red (higher expression) and blue (lower expression). e, Gene ontology GSEA for the top 7 and bottom *7* enriched pathways with normalized enrichment score (NES) on x-axis and descriptive term on y-axis. Dot size represents the number of gene hits in the pathway and dot color represents the – log_10_ p-value. f, CD8^+^ T cell volcano plot with average Log_2_ fold change (x-axis) and –log_10_ p-value (y-axis) for prediabetic vs non-diabetic (reference) PWH. Genes that had ≥ 0.25 log_2_ fold change and adjusted p-value < 0.05 were colored red (higher expression) and blue (lower expression). Abbreviations: NES, normalized enrichment score.

**Figure 7.**
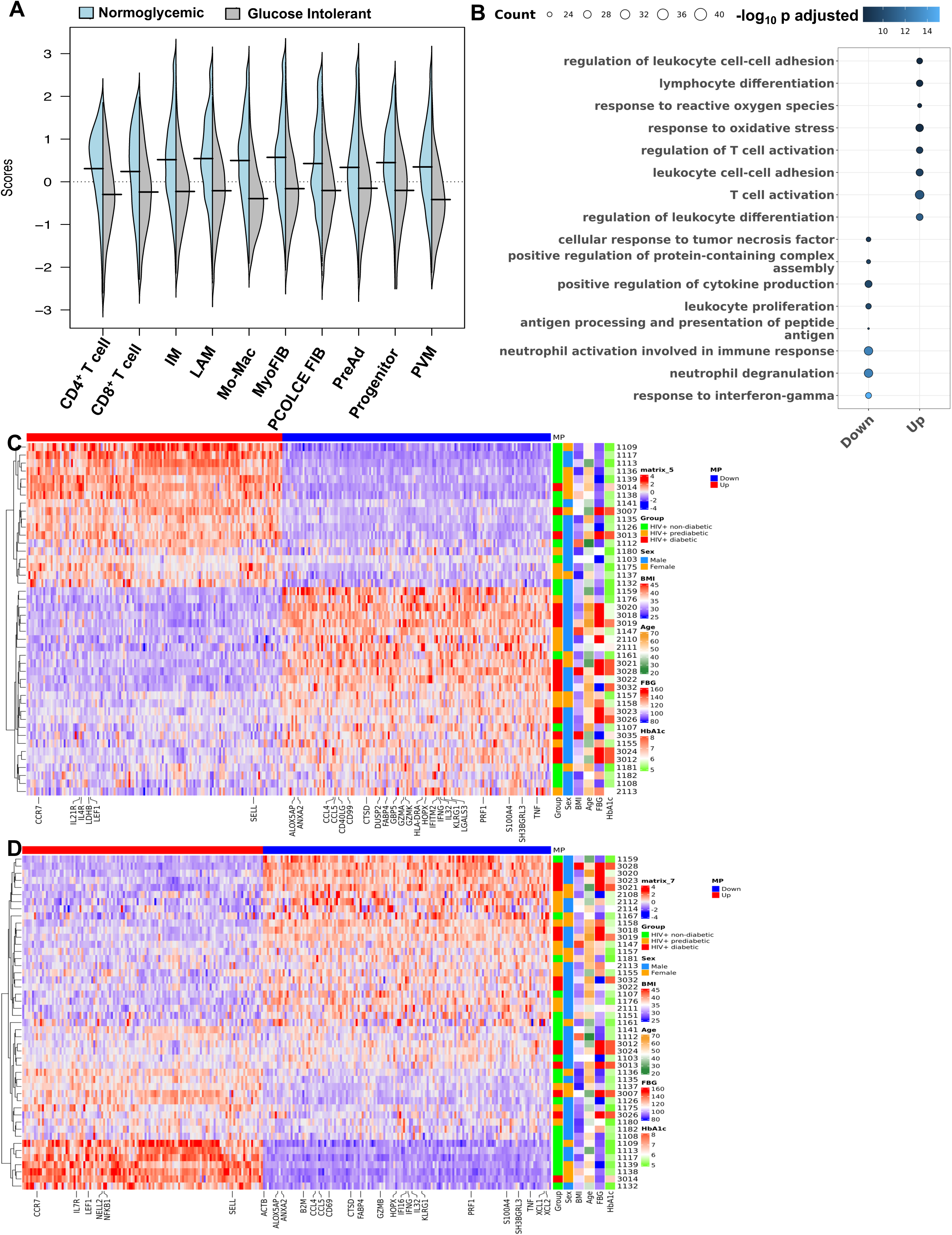
Inter-Cellular Gene Correlation Reveals an Interferon-g, tumor necrosis factor, and lipid metabolism Expression Pattern with Glucose Intolerance. **a**, Distribution of expression scores for each cell type component for multicellular program (MP) 1 with kernel density estimates. b, Over-representation analysis using Gene Ontology for genes in MP1 with higher expression (Up) and lower expression (Down). Dot size represents the number of gene hits in the pathway and dot color represents the –log_10_ p-value. c, Average scaled expression of top CD4^+^ T cell genes from MP1 sorted by expression (columns), across samples plotted with hierarchical clustering (rows) and labeled with clinical variables including body mass index (BMI), age, sex, and measures of glucose intolerance. d, Average scaled expression of top CD8^+^ T cell genes from MP1 sorted by expression (columns), across samples plotted with hierarchical clustering (rows) and labeled with clinical variables including BMI, age, sex, and measures of glucose intolerance. Abbreviations: FIB, fibroblast; IM, intermediate macrophage; Mac, macrophage; Mo, monocyte; LAM, lipid-associated macrophage; PreAd, preadipocyte; PVM, perivascular macrophage.

There were no differentially expressed genes in preadipocytes from prediabetic versus non-diabetic PWH (Supplementary Table 8). Comparison of diabetic with non-diabetic PWH showed similar, but fewer differentially expressed genes between immune cells, which could reflect treatment effect (Supplementary Table 9). Taken together, macrophages show a transcriptional profile that shifts from an immune regulatory function to a metabolic phenotype in prediabetic PWH. Expression of transcription factors associated with M2 polarization are reduced in prediabetic PWH, suggesting decreased differentiation of monocyte-macrophages into PVMs. The T cell transcriptional profile shifts towards an activated effector memory/antigen presentation phenotype in prediabetic PWH. While most of the differentially expressed genes were unique by cell type, there were overlapping genes between cell types in both higher and lower expressed genes, particularly between CD4^+^ and CD8^+^ T cells and between macrophage subsets (Supplementary Fig. 5d, 5e).

### Intercellular Gene Programs Related to Interferon-γ, Tumor Necrosis Factor, and Lipid Metabolic Processes Characterize Glucose Intolerance

Based on some overlapping features of gene changes, and an overall shift in transcriptional profile, we next assessed whether there were coordinated, inter-cellular gene expression programs that define the transcriptional patterns associated with glucose intolerance. We employed DIALOGUE, which is a dimensionality reduction technique that identifies gene expression programs between cell types to identify tissue-specific cellular programs^58^. Multicellular program 1 (MP1) was highly associated with normoglycemia while glucose intolerance was inversely related to MP1 (Fig. 7a). Over-representation analysis of the MP1 showed enrichment of genes related to leukocyte differentiation, regulation of T cell activation, and response to oxidative stress in its upregulated compartment. In contrast, MP1 downregulated compartment was enriched in genes related to IFN-γ responses, neutrophil degranulation, and antigen processing and presentation (Fig. 7b). Glucose intolerance was highly related to CD4^+^ T cell gene expression pattern (Fig. 7c). In CD4^+^ T cells, non-diabetic individuals had increased expression of several genes including *CCR7, LDHB, LEF1, SELL, IL21R* and several ribosomal proteins consistent with a less differentiated state, while prediabetic and diabetic individuals had greater expression of genes related to cytotoxicity (*CCL5, CCL4, PRF1*), activation (*IL32, CD40LG, HLA*), and inflammation (*GBP5, IFNG, TNF, IFITM2)*. Several individuals classified as non-diabetic or prediabetic were borderline based on dichotomous classification, which could explain why some individuals did not cluster as expected (*e.g.* 1182). Several non-diabetic individuals who clustered with glucose-intolerant individuals (*e.g.* 1159, 1108) were cytomegalovirus (CMV) seropositive and had high proportion of cytotoxic T_EMRA_ cells based on flow cytometry, likely in response to CMV infection. This suggests that while glucose intolerance is highly associated with MP1, other factors can contribute to an inflammatory SAT environment. CD8^+^ T cells showed a similar expression pattern; however it did not associate as strongly with glucose intolerance (Fig. 7d). The macrophage compartment MP1 expression program was also not as strongly associated with diabetes status as the T cell compartment but did show enrichment of genes associated with M2 polarization (*KLF10, KLF4, MAFF, MAFB, ATF4, EGR1, EGR2*) and chemotaxis (*CCL2, CCL3, CCL4, CCL8*) in MP1 upregulated compartment, and enrichment of genes associated with lipid metabolism (*APOE, TREM2, APOC1*), and IFN-γ (IFNGR2, IFI27, ISG15) in MP1 downregulated compartment (Supplementary Figures 6a-d). Diabetes status was also significantly associated with MP1 in preadipocytes (Supplementary Figure 6e). Genes associated with non-diabetic phenotype included those important for adipogenesis (*CEBPB, GPX3, KLF2, KLF6, HMOX1, EGR1, EGR2, ZFP36*) whereas those associated with glucose intolerance phenotype included genes related to ECM (*TIMP1, POSTN, COL14A1, PLAAT4, PDGFD*). Overall, MP1 differentiates between normoglycemic individuals with a program enriched for genes related to naïve T cells, immunoregulatory macrophage function, and adipogenesis, as compared to glucose intolerant individuals with a program enriched for genes related to cytotoxicity, inflammation (*TNF*, *IFNG*), cholesterol and lipid metabolism, and ECM deposition.

### CD4^+^ T_EM_ Expressing CD69 are Associated with Changes in Mature Adipocyte Gene Expression

CD4^+^ T_EM_ proportion and transcriptional expression were strongly associated with glucose intolerance compared with other immune cell types. These cells express CD69, which is often a marker of tissue residency^47^. Given the close association with glucose intolerance and MP1, we next assessed whether these cells are associated with changes in adipocyte gene expression patterns. We performed probe-based RNA transcript quantification of whole adipose tissue biopsies for 77 adipocyte genes using the NanoString platform. We found that both the cytometric sorted CD4^+^CD69^+^ T cells and single-cell CD4^+^ T_EM_ cell proportions were associated with higher expression of *ADIPOQ, LPL,* and *LEP,* and lower expression of genes related to long-chain fatty acid metabolism (*CPT1B, CYP27A1, SLC27A5, ACAA1*) and GLP1R in whole SAT (Supplementary Table 10). No other immune cell subsets, including macrophage types or CD8^+^ T_EM_ had a significant relationship with adipocyte gene expression. Taken together, changes in SAT CD4^+^ tissue resident cell composition are associated with changes in adipocyte gene expression, which suggests a potential mechanistic link between SAT immune cells, mature adipocytes, and development of metabolic phenotypes.

### Regulatory Programs Associated with Diabetes in PWH are Also Present in HIV-negative Persons with Diabetes

Finally, we examined whether the regulatory patterns that define glucose intolerance in PWH are similar in diabetic HIV-negative persons. To this end, we recruited 32 diabetic HIV-negative persons with clinical and demographics features similar to the diabetic PWH, and we performed analyses on an integrated dataset of these two groups to assess for differences by HIV status (Supplementary Table 11). We recovered the same cell types in HIV-negative persons observed in the PWH. Compared with the cell distribution of diabetic PWH, HIV-negative persons had fewer lymphoid cells as a proportion of all cell types (Fig. 8a), fewer CD4^+^ cytotoxic T cells (Fig. 8b), but similar macrophage polarization, fibro-adipogenic cell proportions, and CD8^+^ T cell distribution. Transcriptionally, T cells and macrophages from diabetic PWH and diabetic HIV-negative persons were similar (Supplementary Table 12).

**Figure 8.**
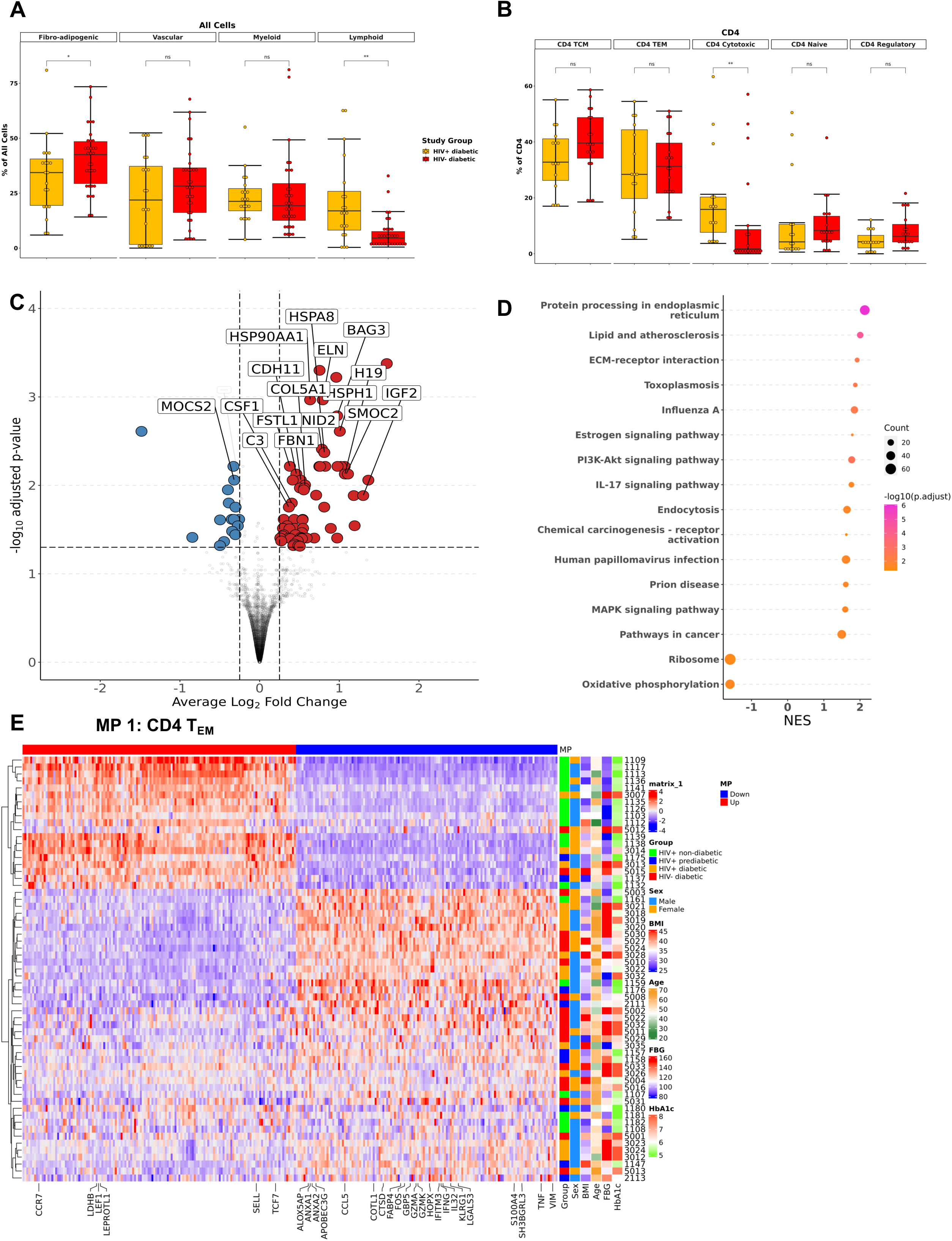
HIV-negative diabetic and HIV-positive diabetic Have Similar Macrophage and T Effector Memory Cell Polarization and Preserved Inter-cellular Gene Expression Program. **a**, Boxplot showing the proportion of major cell categories (stromal, vascular, lymphoid, and myeloid) as a percentage of total cells split by disease status (n = 51). b, Boxplot showing the proportion of macrophage subsets as a percentage of total macrophage cells split by disease status (n = 51). c, Preadipocyte volcano plot with average Log_2_ fold change (x-axis) and –log_10_ p-value (y-axis) for HIV-positive diabetic vs HIV-diabetic (reference) persons. Genes that had ≥ 0.25 log_2_ fold change and adjusted p-value < 0.05 were colored red (higher expression) and blue (lower expression). d, Gene set enrichment analysis (GSEA) using the Gene Ontology database. The top 6 and bottom 6 enriched pathways were included with normalized enrichment score (NES) on x-axis and descriptive term on y-axis. Dot size represents the number of gene hits in the pathway and dot color represents the –log_10_ p-value. e, Average scaled expression of top CD4^+^ T cell genes from MP1 sorted by expression (columns), across samples plotted with hierarchical clustering (rows) and labeled with clinical variables including body mass index (BMI), age, sex, and measures of glucose intolerance. Abbreviations: BMI, body mass index; FBG, fasting blood glucose; HbA1c, hemoglobin A1c; IM, intermediate macrophage; Mac, macrophage; Mo, monocyte; LAM, lipid-associated macrophage; PVM, perivascular macrophage

However, the preadipocyte cells were strikingly different with higher expression of gene related to ECM-, genes that impair adipogenesis, and lipid-processing genes in PWH, suggesting impaired adipogenesis and ECM deposition in PWH compared with HIV-negative (Fig. 8c, 8d, Supplementary Table 12). Finally, with the addition of diabetic HIV-negative persons to the intracellular gene correlation, they tended to cluster with glucose intolerant PWH, suggesting a similar pattern of adipose tissue regulation (Fig. 8e). Overall, we find that although HIV induces significant changes to the lymphoid compartment, the pattern of dysregulation in the adipose tissue of diabetics is similar irrespective of HIV status and our findings may be generalizable to the broader population.

## Discussion

In this study, we generated a comprehensive single cell molecular atlas of SAT in PWH to uncover compositional and transcriptional patterns that are associated with glucose intolerance. We showed a shift towards LAM and LAM-like macrophages and T_EM_ cells. Transcriptionally, macrophages shifted from an immunoregulatory cytokine profile towards a lipid processing phenotype while CD4^+^ and CD8^+^ T cells shifted towards differentiated and effector phenotypes. Intercellular correlation of gene expression demonstrated upregulation of IFN-γ and TNF-related pathways in CD4^+^ and CD8^+^ T cells, upregulation of lipid-processing genes in macrophages, and increased expression of fibrotic genes in preadipocytes in PWH with glucose intolerance. CD4^+^ memory cells expressing CD69 were most strongly associated with glucose intolerance and alterations in adipocyte gene expression, providing a plausible mechanistic link to altered mature adipocyte function and development of metabolic syndrome. Finally, we show that this regulatory pattern is preserved in HIV-negative participants with diabetes. These data suggest our findings may be generalizable and should be further investigated in HIV-negative cohorts.

Our characterization of SAT resident immune cells and fibro-adipogenic precursor cells expand on previous scRNA-seq descriptions of human adipose tissue in HIV-negative persons^14, 15, 23^. Uniquely, we evaluated the role of glucose intolerance in compositional and transcriptional polarization of adipose tissue and leveraged our large dataset to evaluate the independent contributions of important biological factors including sex, age, and BMI. We showed a coordinated gene expression response that skews towards inflammatory and lipid metabolism phenotype with glucose intolerance, which may perpetuate ongoing adipose tissue inflammation. While we chose scRNA-seq to perform CITE-seq in parallel, current methods for making single-cell suspensions for scRNA-seq do not capture mature adipocytes. Therefore, we used whole tissue to capture adipocyte gene expression and demonstrate that CD4^+^ T cells expressing CD69, a marker of tissue residency and a group comprised primarily of T_EM_, have significant association with the adipocyte transcriptional profile. This could provide a biological basis linking the SAT immune cell changes with overall metabolic health.

Our study had some limitations. The cross-sectional design precluded an assessment of the temporal course of compositional and transcriptional changes, and future longitudinal scRNA-seq studies are needed. Our separation of participants into three groups by glucose intolerance meant some individuals were on the margin between states, though this was addressed, in part, by our analysis of FBG as a continuous endpoint. Additionally, our cohort lacked a non-diabetic or pre-diabetic HIV-negative group. Despite the size of our unbiased molecular atlas of SAT, we may have missed low frequency cells that contribute to inflammation. Finally, it is difficult in clinical studies to account for potential confounders. However, we overcame this with reasonably matched groups and a large overall cohort to model the contributions of important biological factors to composition and transcriptional patterns.

In summary, we found unique SAT compositional and transcriptional changes with glucose intolerance and identified a conserved cellular regulatory program that differentiated non-diabetic and glucose intolerant individuals. Our dataset is publicly available to the research community on an interactive platform (http://vimrg.app.vumc.org/). These data provide insight into the complexity and breadth of SAT cells that may contribute to glucose intolerance and accelerate future investigation into the role of fibro-adipogenic and immune cell interactions that may open new avenues of research and lead to developing therapeutic interventions.

## Methods

### Study Participants

Participants were members of the HIV, Adipose Tissue Immunology, and Metabolism (HATIM) study developed to evaluate adipose tissue characteristics in the context of HIV infection and metabolic disease (ClinicalTrials.gov registration NCT04451980). PWH were recruited from the Vanderbilt University Medical Center Comprehensive Care Clinic between August 2017 and June 2018, were on antiretroviral therapy (ART) for ≥ 18 months, had virologic suppression (serum HIV-1 RNA quantification < 50 copies/mL) for ≥ 12 months, had a CD4^+^ T cell count ≥ 350 cells/mm^3^ and had no known inflammatory or rheumatologic conditions. Participants were classified as non-diabetic (HbA1c < 5.7% and/or fasting blood glucose [FBG] < 100 mg/dL), pre-diabetic (HbA1c 5.7-6.5% and/or FBG 100-125 mg/dL), or diabetic (HbA1c > 6.4% and/or FBG ≥ 126 mg/dL and/or on anti-diabetic medication) in accordance with the American Diabetes Association criteria ^59^. HIV-negative participants with diabetes were simultaneously recruited from the Vanderbilt *ResearchMatch* cohort, and these individuals were group matched by age and BMI with diabetic PWH. All participants underwent a single clinical research visit after a minimum 8-hour fast that included HbA1c and FBG measurement, peripheral blood mononuclear cell (PBMC) collection, fasting plasma collection, and anthropomorphic measurements. All individuals also underwent subcutaneous adipose tissue liposuction as described below. The Vanderbilt Institutional Review Board approved the research, and all participants provided written consent.

### Adipose Tissue Collection and Processing

Subcutaneous adipose tissue biopsies were collected approximately 3 centimeters to the right of the umbilicus after anesthetizing the skin with lidocaine/epinephrine and infiltrating 40 mL of sterile saline and lidocaine into the SAT. We collected approximately 5 grams of adipose tissue using a 2.1 mm blunt, side-ported liposuction catheter (Tulip CellFriendly**^TM^**GEMS system Miller Harvester, Tulip Medical Products) designed for extraction of viable adipocytes and stromal vascular fraction (SVF) during cosmetic adipose tissue transfer procedures^60^. Using this method, adipose tissue is recovered in droplets generally < 3 mm in diameter, limiting the need for mechanical dissociation. The tissue was placed in 40-50 mm^3^ of cold saline and mixed. Visible blood clots were removed, and the sample was transferred to a 70 µm filter for repeat saline washes with constant stirring. The adipose tissue was then placed in a gentleMACS**^TM^** Dissociator (Miltenyi Biotec) followed by incubation with 100 µL of collagenase D (20mg/mL). The SVF was separated using a Ficoll-Paque Plus density gradient. Samples were cryopreserved in fetal bovine serum with 10% DMSO in liquid nitrogen.

### Library Preparation and Sequencing

An antibody Total Seq C master mix containing 45 common markers for lineage, memory, and activation was created by adding 0.5 µL of each antibody. Twelve samples, with representative samples from the four metabolic groups (non-diabetic PWH, pre-diabetic and diabetic PWH, and HIV-negative diabetics), were processed at a single time. The samples were quickly thawed and transferred to labeled 15 mL tubes and diluted to 10 ml with phosphate-buffered saline (PBS) before spinning down at 300 G for 10 minutes. The supernatant was aspirated, and the pellet was resuspended in cell staining buffer and transferred to labeled flow tubes. The sample was again spun at 300 G for 10 minutes, and the supernatant was aspirated. Cells were resuspended in 100 µL of staining buffer. Human TruStain FcX Receptor Blocking Solution (5 µL) was added to the tube and incubated at 4 Celsius for 10 minutes. The antibody master mix was spun at 15000 rpm for 5 minutes to remove any aggregates that had formed, and 22.5 µL of the antibody mix was added to each tube of cells and mixed by flicking the tube. A total of one µL of Total Seq C hashtag antibody was added to the samples, and the samples were incubated at 4 Celsius for 30 minutes. After incubation, the cells were washed three times with Cell Staining Buffer and spun at 300 G for 10 minutes. The cells were resuspended in around 100 µL of PBS with 0.04% bovine serum albumin (BSA). The cells were counted using the Countess II Automated Cell Counter (Thermo Fisher) to determine the suspension volume to transfer to obtain 5,000 cells. Four samples (each with a unique hashtag antibody) were pooled by metabolic status (except non-diabetic and prediabetic samples from PWH which were pooled) together into one tube. The multiplexed single cells were loaded onto a Chromium Single Cell 5’ v.2 assay (10x Genomics). Libraries were sequenced on the NovaSeq 6000 S2 platform (Illumina). Illumina bcl files were demultiplexed using bcl2fastq. Raw reads were then aligned to the human genome (hg38) using STAR v. 2.7.2a, and cells were called using Cellranger count (v.6.0.0) with default settings. Souporcell, which leverages single nucleotide variants to assign individual cells to genotypes and generate a VCF file, was used to genetically demultiplex the samples^61^. SoupX was used with default parameters to remove ambient RNA contamination from the count matrice^62^.

### Quality Control

The R Statistical Programming package Seurat V4 was used to further process the scRNA-seq data^63^. First, cells with > 25% mitochondrial gene expression, < 800 transcript reads, and < 200 genes were filtered out. The threshold of 800 transcript reads per cell was selected because the performance of Souporcell begins to decrease with fewer transcripts^61^. Hashtag oligonucleotides (HTOs) were normalized using centered log-ratio (CLR) transformation, and cells were assigned HTO implemented in the function HTODemux with default parameters^25, 64^. We then added the Souporcell assignment to the metadata and linked the Souporcell cluster designation with the HTO classification, forming a link with the metadata. All cells that were unassigned by Souporcell were removed. To identify heterotypic doublets, we used standard processing of each lane, including normalization and variance stabilization using regularized negative binomial regression (SCTransform), dimensional reduction, and clustering^65^. DoubletFinder^66^, was used to identify potential heterotypic doublets in the data using Souporcell designation as the ground truth. Cells that were identified as doublets by Souporcell or DoubletFinder, clusters where > 60% of cells were identified as doublets, and clusters that expressed transcripts from multiple major lineages (endothelial, immune, fibro-adipogenic, vascular), were removed after integrating the datasets.

### Cell Clustering and Annotation

Processed individual lanes were merged using the merge function. The gene counts were normalized for each cell by dividing by the total gene counts and multiplying by a factor of 10,000 before applying log-transformation as implemented by the LogNormalize function. Protein expression counts were normalized using the CLR method. Variable genes were identified using the FindVariableFeatures function (nFeatures = 3000), and the data were scaled and centered using the ScaleData function. Principal component analysis (PCA) was performed on the scaled data. The number of principal components (PCs) used for downstream analysis was selected based on elbow plots and heat maps of PC dimensions as implemented in the DimHeatmap function. To reduce the batch effect associated with running multiple 10X lanes, we used the Harmony algorithm on the uncorrected PCs implemented as a Seurat wrapper to integrate across lanes^26^. To evaluate the effectiveness of integration, we used the SCIB pipeline^67^. The overall metric ranking was calculated by the summation of the scaled overall bio-conservation score * 0.6 + batch score * 0.4. We performed clustering on the Harmony-corrected PCs using FindNeighbors and FindClusters functions. Marker genes for each cluster were determined using the Wilcoxon Rank Sum test implemented in FindAllMarkers.^68^ The integrated dataset was then subclustered as described below. After processing each subcluster, they were merged again, and the process described above (except normalization) was repeated to obtain a cell atlas. We performed manual annotation of cell populations based on canonical markers and markers previously identified in scRNA-seq (Supplementary Table 4).

### Subclustering Analysis

We subclustered on major cell types, including fibro-adipogenic (*COL1A2, CCDC80*), vascular (*CLDN5, ACTA2*), myeloid (*LYZ, CD68, CD14, CD1C, LILRA4, CLEC9A*), and lymphoid (*CD3, NKG7*). We followed the procedure described previously for each subcluster but did not repeat RNA and protein normalization. Clusters defined by mitochondrial gene expression and/or transcriptional doublets were removed from the analysis. We performed differential gene expression using the FindAllMarkers function implemented in Seurat to identify gene markers for each cell population and included genes that were expressed in 25% or more cells and had a log_2_ fold change of 0.25 or greater.

### Composition Analysis

Individuals contributing < 30 cells in the subset investigated were excluded. We compared cell composition between disease states for each subcluster by evaluating each cell type as a proportion of total cells in the subcluster. We assessed whether cell type changes were significant between non-diabetic PWH and prediabetic PWH, and between non-diabetic PWH and diabetic PWH using the Wilcoxon Rank Sums test. P values were adjusted for multiple comparisons using Benjamini-Hochberg procedure.

To evaluate the independent relationship of cell proportions with BMI, age, and measures of glucose intolerance (HbA1c, FBG), we used partial spearman’s correlation implemented in PResiduals^69^. We visualized the results by plotting spearman’s ρ per cluster superimposed on the uniform manifold approximation and projection (UMAP). We used the same method to evaluate the inter-cellular proportion relationships. To evaluate the independent relationship of female sex with cell proportions, we used an ordinal linear regression with cell proportion as the outcome and sex (male reference) as the independent variable adjusted for age, BMI, and diabetes status. The β coefficient was converted to an odds ratio (female: male) going from the 25^th^ to 75^th^ percentile (proportion) and visualized by plotting on the UMAP as above.

### Transcriptional Analysis

To evaluate differentially expressed genes between non-diabetic and pre-diabetic PWH, we aggregated gene counts (psuedobulk) for each individual using scuttle^70^. We then used a negative binomial generalized linear model implemented in DESeq2 to evaluate differentially expressed genes adjusting for age, sex, and BMI. We used clusterProfiler to perform gene set enrichment analysis using the Kyoto Encyclopedia of Genes and Genomes (KEGG) and Gene Ontology (GO) biological processes to identify pathways that were enriched in prediabetic compared with non-diabetic PWH.^71^

### Pseudotime Analysis

The R package Slingshot,^56^ was used to assess the pseudo-time trajectory of PVM. Monocyte-macrophages were specified as the root cluster with the dimensionality reduction produced by UMAP. To identify temporally dynamic genes, we fit a generalized additive model (GAM) using a negative binomial additive model implemented in the R package tradeSeq^57^. We used the associationTest function with l2fc set to 2 to identify genes significantly changing along the pseudo-time, defined as FDR-adjusted p-value < 0.05. The genes were then ordered according to pseudo-time and plotted by scaled expression using ComplexHeatmap^72^.

### Multicellular Gene Expression Programs

To evaluate for a coordinate cellular program that characterizes glucose intolerance, we used DIALOGUE^58^, evaluating the cell types we previously identified as associated with glucose intolerance including CD4^+^ and CD8^+^ T cells, macrophage subsets, preadipocytes, and fibroblast cells. DIALOGUE was run with default settings. The multi-level models were fit by glucose intolerance status, and adjusted for technical variability, sex, age, and BMI. Average scaled expression of top genes from MP1 were sorted by expression and samples were plotted with hierarchical clustering (rows) and labeled with clinical variables including BMI, age, sex, and measures of glucose intolerance using ComplexHeatmap.

### Whole Adipose Tissue Messenger RNA Expression

The detailed methods for these samples have been published elsewhere^73^. Briefly, messenger RNA (mRNA) was extracted from cryopreserved SAT with Qiagen RNeasy Lipid Tissue Kit after mechanical lysis. The NanoString nCounter Plex platform (NanoString, Seattle, WA) was used to quantify mRNA transcripts for 77 genes related to adipocyte function. The mRNA count was normalized using eight synthetic spike-ins for negative control and six synthetic spike-ins for positive controls. The coefficient of variation (CV) was calculated for control genes. First, the mean of the negative controls was used as the background level and subtracted from each gene count. The normalization factor for the mRNA content was calculated using the geometric mean of a set of pre-specific housekeeping genes. The count data were then divided by the normalization factor to generate counts normalized to the geometric mean of housekeeping genes. The normalized gene count data was then log2-transformed, and a linear regression model was used to assess the relationship between the cell-type proportion with mRNA expression. The cell proportion was the independent variable, and the log2-transformed mRNA count was the dependent variable, adjusted for age, sex, BMI, metabolic status, and batch. Benjamini-Hochberg was used to correct for multiple comparisons.

### Data Availability

Raw sequencing files, cell metadata, and fully integrated Seurat objects are available via the NCBI GEO with the primary accession code GSE198809. The data is available on an interactive ShinyCell-based^74^, web application at https://vimrg.app.vumc.org/.

### Code Availability

All code processing and analyses code for this paper are available on github at https://github.com/VIMRG/AdiposeTissueAtlas.

## Acknowledgement

This work was supported by National Institutes of Health grants R01DK112262 (JRK and CNW), R01HL145372 (JAK), K23HL156759 (CNW, JRK), the Vanderbilt Clinical and Translational Science Award from NCRR/NIH grant UL1RR024975, the Vanderbilt Infection Pathogenesis and Epidemiology Research Training Program (VIPER) grant T32AI007474, the Vanderbilt Scholars in HIV and Heart, Lung, Blood and Sleep Research (V-SCHoLARS) grant K12HL143956, the Vanderbilt Institute for Clinical and Translational Research supported by grant KL2TR002245 from the National Center for Advancing Translational Sciences, the Tennessee Center for AIDS Research grant P30AI110527 (SAM, CNW, JRK, SAK), the Doris Duke Charitable Foundation grant 2021193 (CNW, JRK) and the Burroughs Welcome Fund grant 1021480 (CNW, JRK). The funding authorities had no role in study design, data collection, analysis, interpretation, the decision to publish, or manuscript preparation. We also thank the participants of the HATIM study. The figures were created with BioRender.com.

## Author Contributions

Conceptualization, C.N.W., J.R.K., S.A.M.; Methodology, C.N.W., J.R.K., S.A.M., R.D.G., J.D.S., A.C., R.R., C.W., J.A.K., S.S.B.; Formal Analysis, S.S.B., C.N.W., J.R.K., S.A.K., J.A.K., R.R., F.Y., R.F.; Investigation, C.N.W., C.W., J.D.S., R.D.G., L.H.; Writing – Original Draft, S.S.B., C.N.W., J.R.K., J.A.K.; Writing – Review & Editing, S.S.B., C.N.W., J.A.K., J.R.K., S.A.M., S.A.K., C.L.G., M.M., L.H., J.D.S., C.W., R.F., F.Y.; Funding Acquisition, C.N.W., J.R.K., S.A.M.

## Declaration of Interests

J.A.K. consults for Boehringer Ingelheim, Janssen, and Bristol-Myers-Squibb, serves on the scientific advisory board for APIE Therapeutics, and provides non-financial study support to Genentech. J.R.K. has served as a consultant to Gilead Sciences, Merck, ViiV Healthcare, Theratechnologies, and Janssen, and has received research support from Gilead Sciences and Merck. C.N.W. serves on the scientific advisory board for ViiV Healthcare.

## Supplementary Figures

**Supplementary Figure 1.**
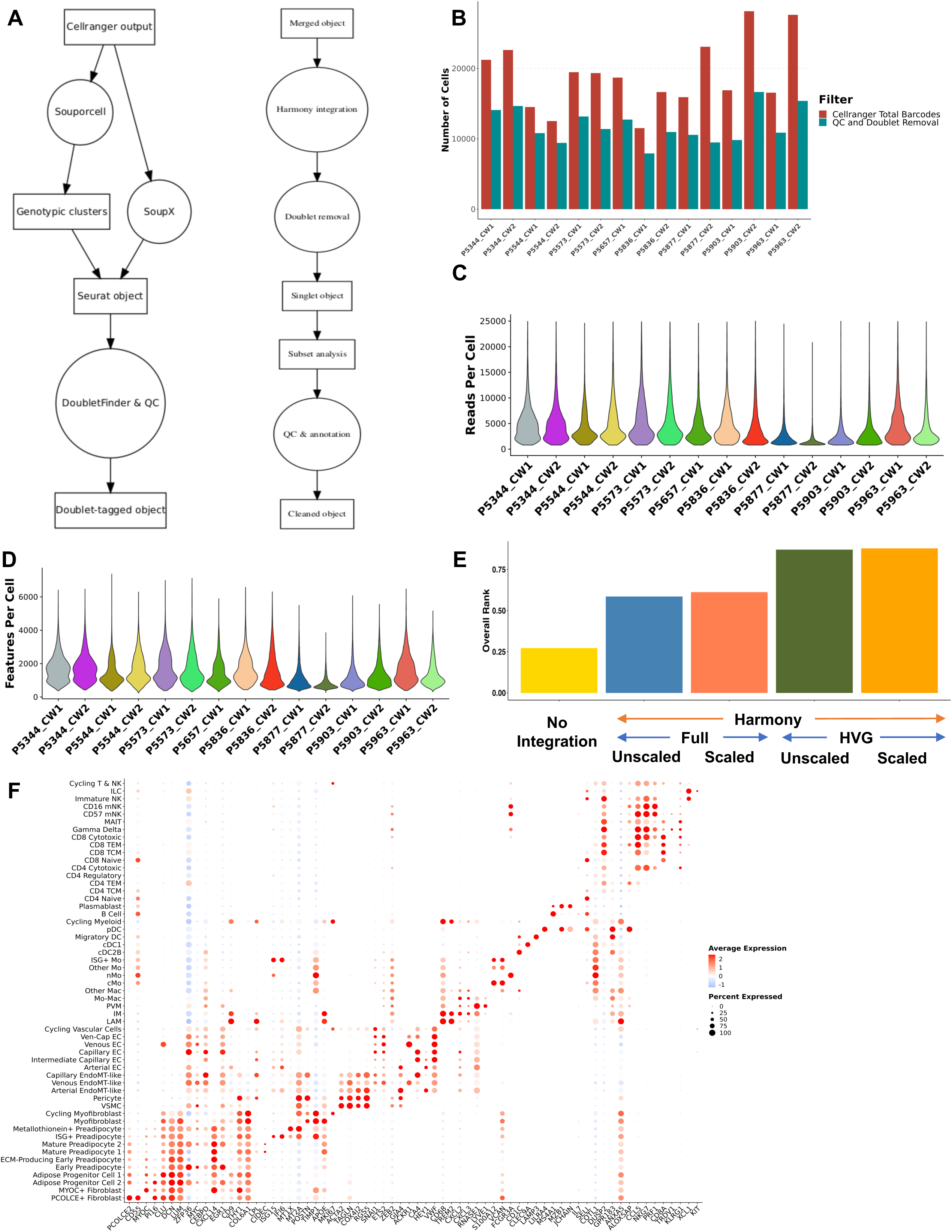
Bioinformatic Pipeline and Quality Metrics. **a**, Flow diagram of bioinformatic pipeline. Cells were called using the Cell Ranger count function with default settings. Soupx was used to remove ambient RNA and the corrected count matrices were used to create a Seurat object. Simultaneously, Souporcell was used to genetically demultiplex the samples. Doublets identified by Souporcell were used as ground truths for DoubletFinder. Next, each Seurat object underwent quality control filtering and downstream dimensional reduction and clustering. The processed objects were then integrated using Harmony correction on the principal components prior to clustering. Cells identified as doublets, clusters with > 60% doublets, or expression of 2 or more major lineages were removed. Cells were subclustered based on major cell lineages and transcriptional doublets were iteratively removed prior to reintegration. **b,** Barplot graph showing each lane (x-axis) and number of cells (y-axis). The graph is split by the total barcodes identified by Cell Ranger (red) and the cells remaining after doublet and QC removal (green). **c,** Distribution of read counts per cell (y-axis) by lane (x-axis). **d,** Distribution of the number of features (genes) per cell (y-axis) by lane (x-axis). **e,** Overall integration metric ranking generated by the SCIB pipeline. Unintegrated, harmony integration with feature selection (HVG) and no feature selection, as well as scaled and unscaled were input in the SCIB pipeline. The overall rank was computed by overall batch score (0.5) and bioconservation (0.6) that were scaled for comparison. **f,** Dot plot with selected genes on the x axis and cell type on the y axis. The size of the dot represents the percentage of cells with expression of that gene and the color of the dot represents the average expression level for that gene.

**Supplementary Figure 2.**
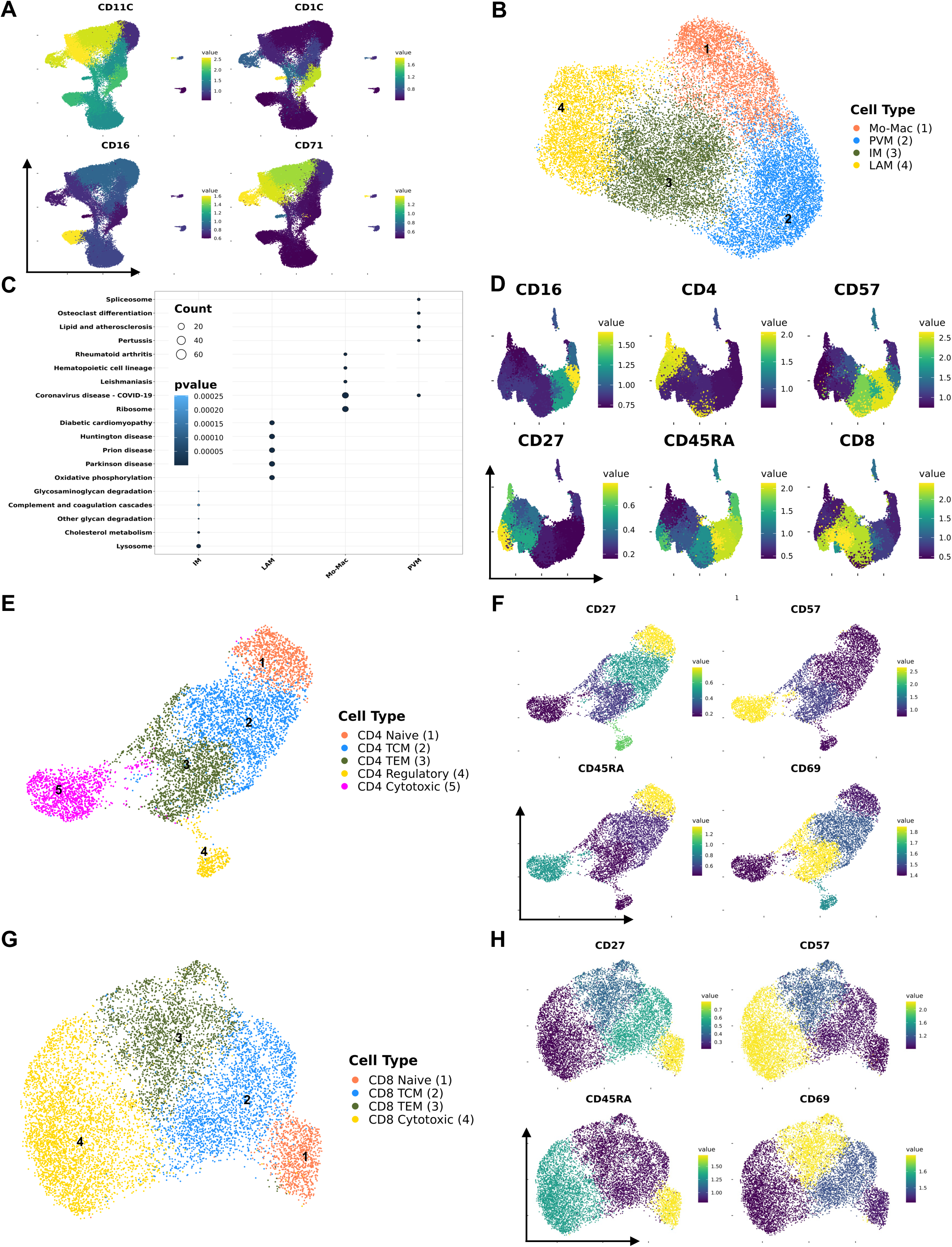
CITE-seq and Pathway Analysis Support Cell Annotation. **a**, Antibody derived tags (ADT) for CD11C, CD1C, CD16, and CD71 plotted as the average cluster expression onto the uniform manifold approximation and projection (UMAP) for myeloid cells. **b**, UMAP of IM, PVM, Mo-Mac, and LAMs from 59 individuals. **c,** Gene Ontology (GO) over-representation analysis with macrophage subtype on the x-axis and GO pathway on the y-axis. The size of the circle denotes the number of genes assigned to the pathway while the color denotes the p-value. **d**, ADT for CD16, CD27, CD4, CD45RA, CD57, and CD8, plotted as the per cluster average expression superimposed on the lymphoid UMAP. **e**, UMAP of CD4^+^ T cells (n = 8,435 cells) subset from all T cells. **f**, ADT for CD27, CD45RA, CD57, and CD69, plotted as the per cluster average expression superimposed on the CD4^+^ T cell UMAP. **g**, UMAP of CD8^+^ T cells (n = 11,356 cells) subset from all T cells. **h**, ADT for CD27, CD45RA, CD57, and CD69, plotted as the per cluster average expression superimposed on the CD8^+^ T cell UMAP. Abbreviations: cMo, classical monocyte; cDC1, conventional dendritic cell type 1; cDC2B, conventional dendritic cell type 2B; DC, dendritic cell; ECM, extracellular matrix; ILC, innate lymphoid cell; IM, intermediate macrophage; IM, intermediate macrophage; LAM, lipid-associated macrophage; Mac, macrophage; mNK, mature natural killer; Mo, monocyte; NK, natural killer; nMo, non-classical monocyte; pDC, plasmacytoid dendritic cell; PVM, perivascular macrophage

**Supplementary Figure 3.**
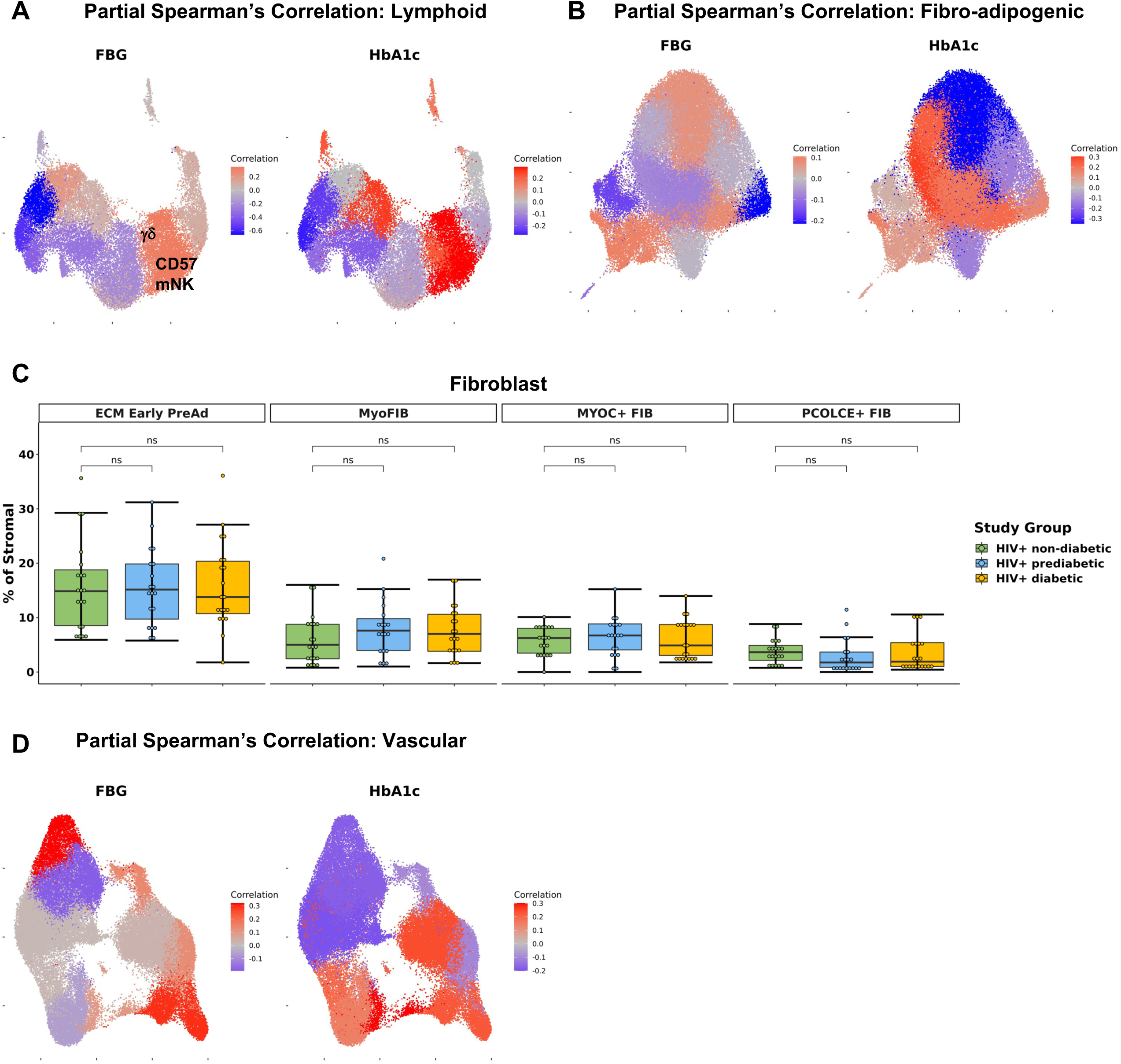
Cell Type Proportions by Diabetes Status. **a**, Partial spearman’s correlations between fasting blood glucose (FBG) or hemoglobin A1c (HbA1c) and lymphoid cell proportions. Spearman’s r for the biological factor (FBG or HbA1c) and each cluster proportion was calculated. The r value (blue, negative; red, positive) for each cluster was plotted onto the macrophage uniform manifold approximation and projection (UMAP). Clusters with significant correlation (p < 0.05) are labeled. **b,** Partial spearman’s correlation between FBG or HbA1c and cluster proportion plotted onto the fibro-adipogenic UMAP. **c**, Boxplot showing the proportion of fibroblast types split by disease state (HIV+ non-diabetic, green; HIV+ prediabetic, blue; HIV+ diabetic, yellow) (n = 59). The horizontal black line represents the median, the box shows the lower and upper quartile limits and the whiskers are 1.5x the interquartile range. **d,** Partial spearman’s correlation between FBG or HbA1c and cluster proportion plotted onto the vascular UMAP. Abbreviations: ECM, extracellular matrix; FIB, fibroblast; mNK, mature natural killer.

**Supplementary Figure 4.**
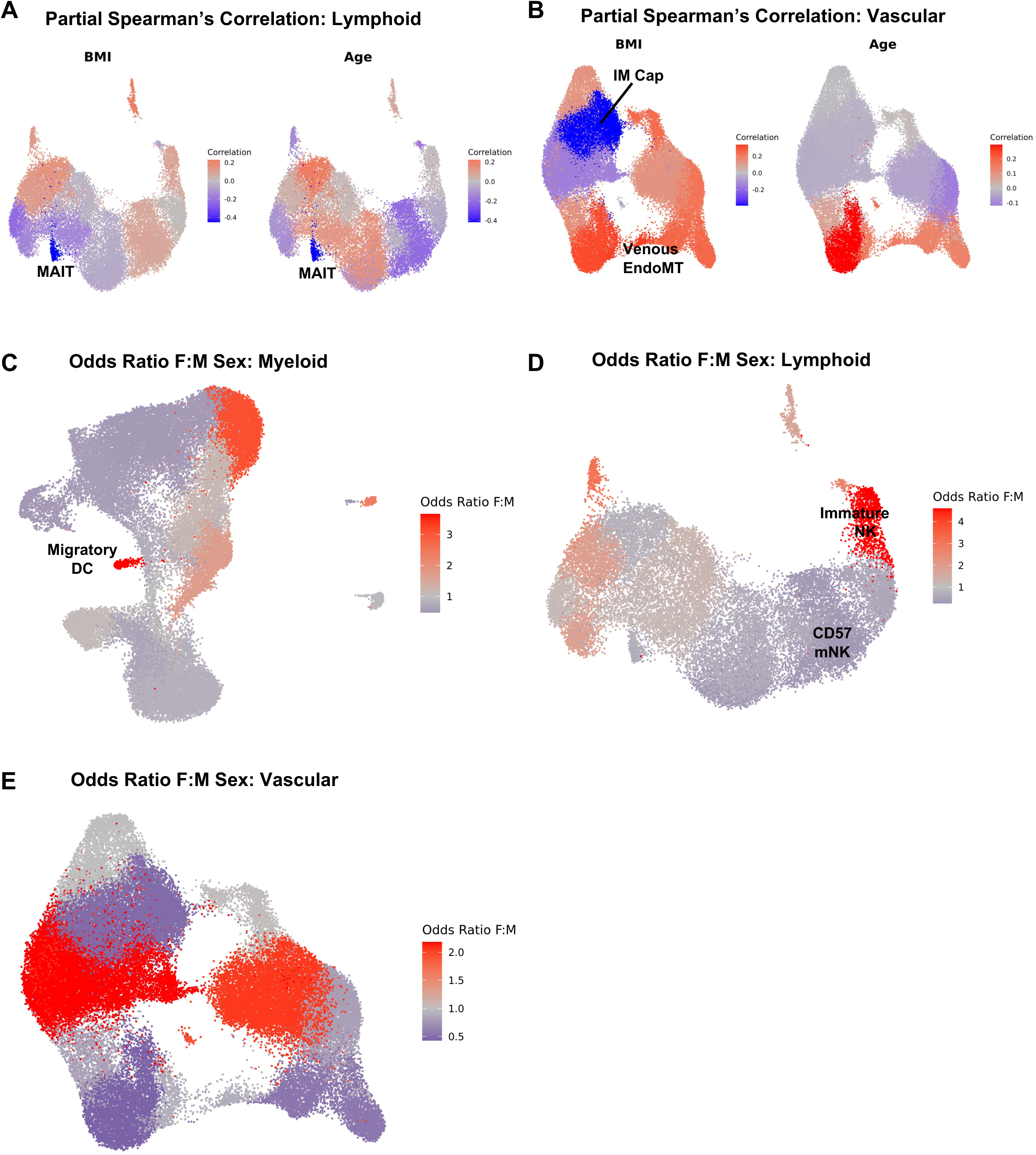
Relationship of Body Mass Index, Age, and Sex to Other Cell Populations. **a-b**, Partial spearman’s correlations. Spearman’s r for the biological factor (body mass index [BMI] or age) and each cluster proportion was calculated. The r value (blue, negative; red, positive) for each cluster was plotted onto the uniform manifold approximation and projection (UMAP). Clusters with significant correlation (p < 0.05) are labeled. BMI and age plotted onto **a,** Lymphoid UMAP, **b,** Vascular UMAP. **c-e,** Ordinal linear regression with cluster proportion as the outcome and sex as the independent variable adjusted for age, BMI, and diabetes status. The regression coefficient for sex was converted into an odds ratio (female: male) and plotted per cluster onto the UMAP. Odds ratio female: male plotted onto **c**, Myeloid UMAP, **d**, Lymphoid UMAP, **f,** Vascular UMAP. Abbreviations: BMI, body mass index; DC, dendritic cell; EndoMT, Endothelial-Mesenchymal Transition; IM Cap, Intermediate Capillary Endothelial;.mNK, mature natural killer.

**Supplementary Figure 5.**
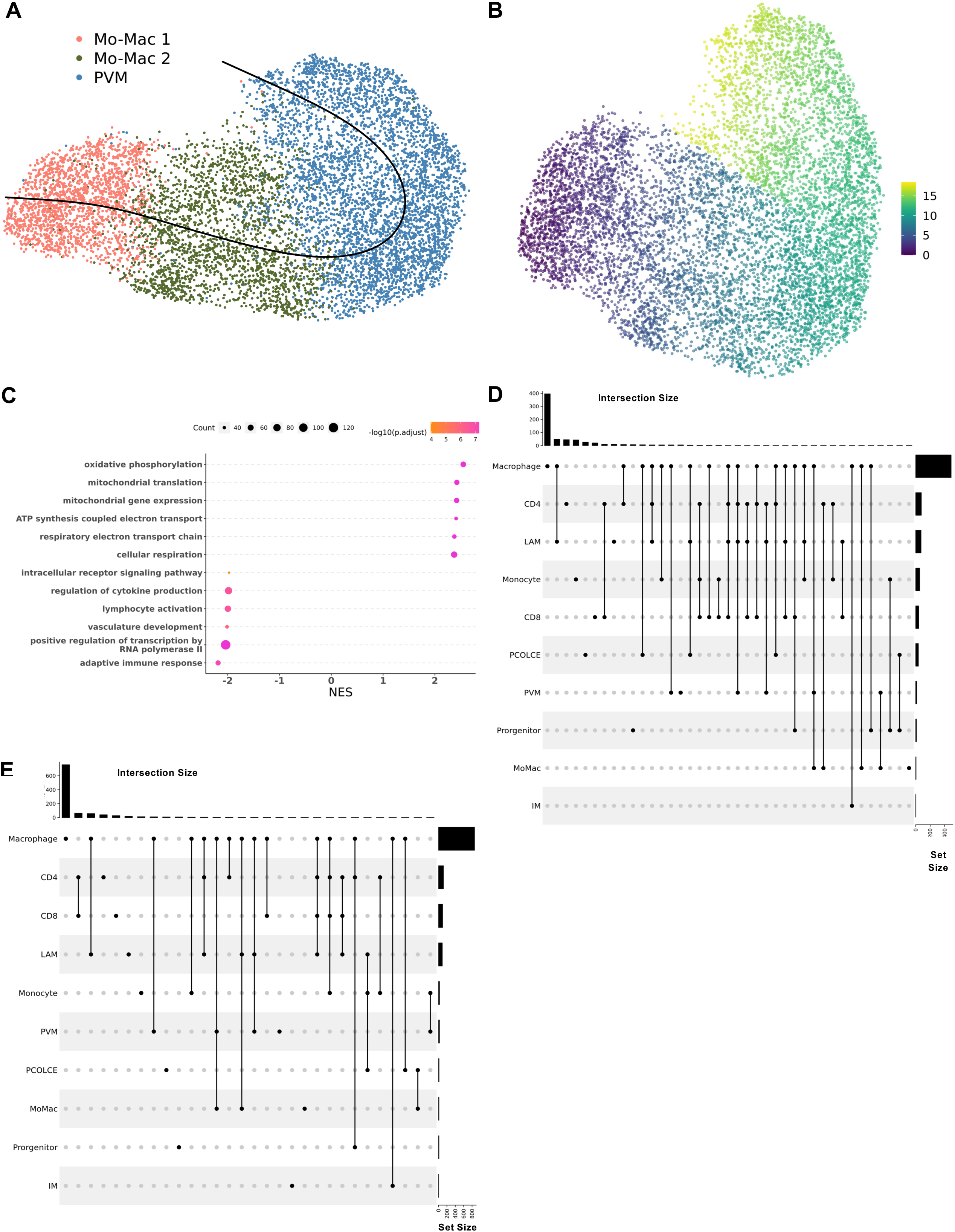
Transcriptional Profile of Adipose Tissue Immune Cells. **a**, Slingshot trajectory with monocyte-macrophage as the root embedded on the UMAP and, **b**, Slingshot pseudotime. **c,** Gene set enrichment analysis (GSEA) using the Gene Ontology database. The top 6 and bottom 6 enriched pathways were included with normalized enrichment score (NES) on x-axis and descriptive term on y-axis. Dot size represents the number of gene hits in the pathway and dot color represents the –log_10_ p-value. **d,** Upset plot for genes with higher expression in prediabetic compared with non-diabetic PWH across cell types. The x-axis shows the set size for each cell type (total number of differentially expressed genes in the cell type) and the y-axis shows the intersection size with other cells (number of shared genes). **e**, Upset plot for genes with lower expression in prediabetic compared with non-diabetic PWH across cell types. The x-axis shows the set size for each cell type (total number of differentially expressed genes in the cell type) and the y-axis shows the intersection size with other cells (number of shared genes). Abbreviations: NES, normalized enrichment score.

**Supplementary Figure 6.**
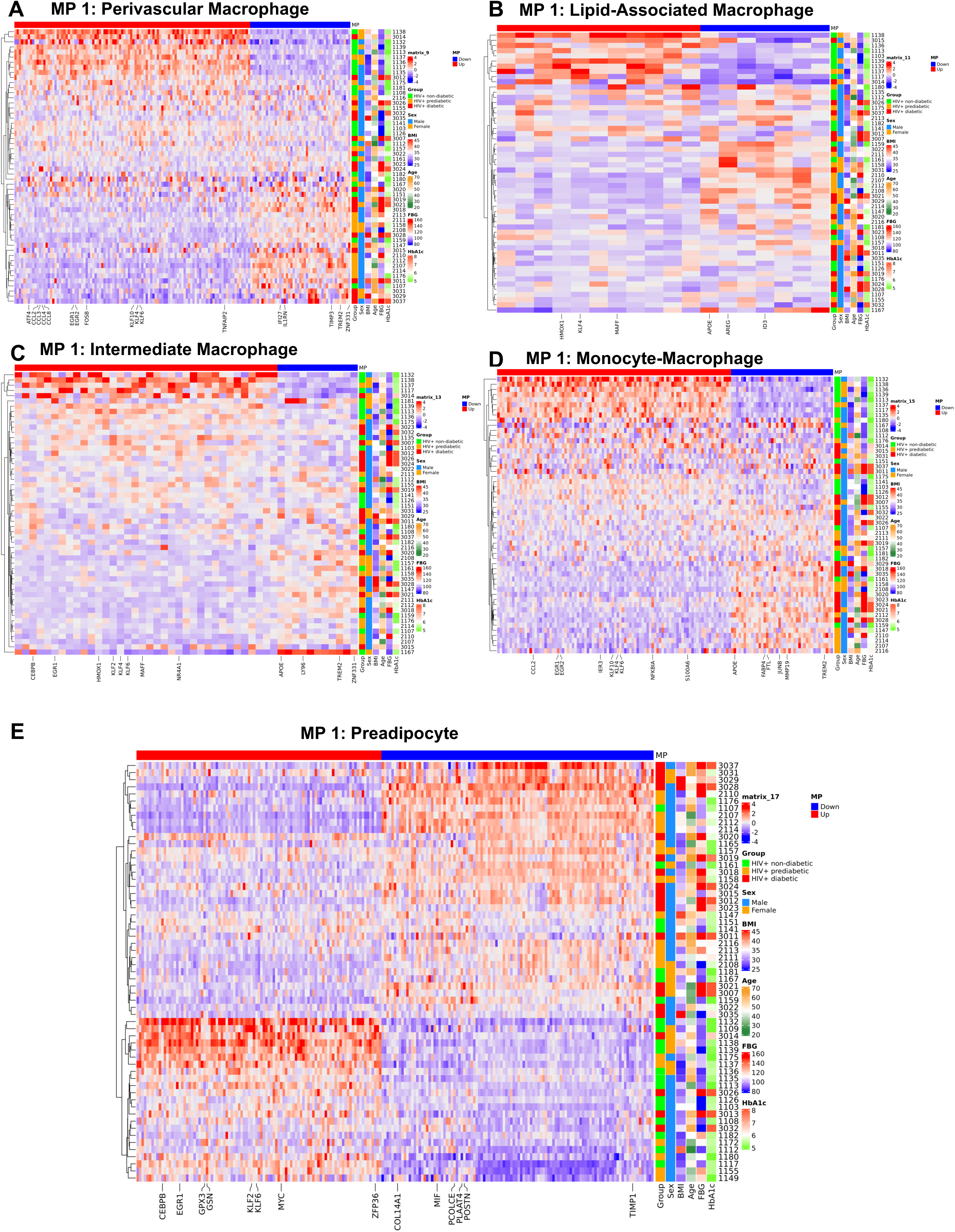
Multicellular Program Using DIALOGUE Identifies Expression Patterns Associated with Glucose Intolerance. **a-f**, Average scaled expression of top genes from MP1 sorted by expression (columns), across samples plotted with hierarchical clustering (rows) and labeled with clinical variables including body mass index (BMI), age, sex, and measures of glucose intolerance. **a,** Perivascular macrophages. **b,** Lipid-associated macrophages. **c,** Intermediate macrophages. **d**, Monocyte-macrophages. **e,** Preadipocytes. Abbreviations: BMI, body mass index; FBG, fasting blood glucose; HbA1c, hemoglobin A1c.

## Notes

https://github.com/VIMRG/AdiposeTissueAtlas

http://vimrg.app.vumc.org/

https://www.ncbi.nlm.nih.gov/geo/query/acc.cgi?acc=GSE198809

### Summary of Updates

Section on transcriptional regulators of glucose intolerance added; figures updated; supplemental files updated.

